# Emergent loss of microbial biodiversity with increasing resource diversity

**DOI:** 10.1101/2025.05.13.653732

**Authors:** Or Shalev, Xiaozhou Ye, Joachim Kilian, Mark Stahl, Christoph Ratzke

## Abstract

The origin of biodiversity is arguably the most fundamental question in ecology — especially how countless microbial species coexist within a single community. A prevailing hypothesis holds that microbial species coexist by specializing on different resources, thereby reducing competition. Accordingly, increasing resource diversity is expected to promote species coexistence and boost biodiversity. Surprisingly, we observe the opposite: microbial biodiversity often declines as resource diversity increases. This unexpected result emerges from a widespread physiological response across diverse microbial taxa, in which more complex environments trigger higher overall resource uptake. This intensifies competition and drives biodiversity loss. Updating a central ecological model to include this physiological trait accurately reproduces the observed decline. Our findings demonstrate that physiological changes at the individual level can overturn classical ecological principles and scale up to restructure entire ecosystems.

## Introduction

Microbial life flourishes in astonishing biodiversity. A single gram of soil can harbor over ten thousand bacterial species^1^. Yet how so many species coexist has remained a central question in ecology for nearly a century^2–4^. Classic theory holds that when species compete for the same resource, the most efficient consumer prevails, driving others to extinction through competitive exclusion^2^. Coexistence, therefore, requires niche differentiation: species avoid direct competition by exploiting different resources^5^. By this logic, greater resource diversity should support more coexisting species^6,7^ (**Fig. 1A**).

**Figure 1:**
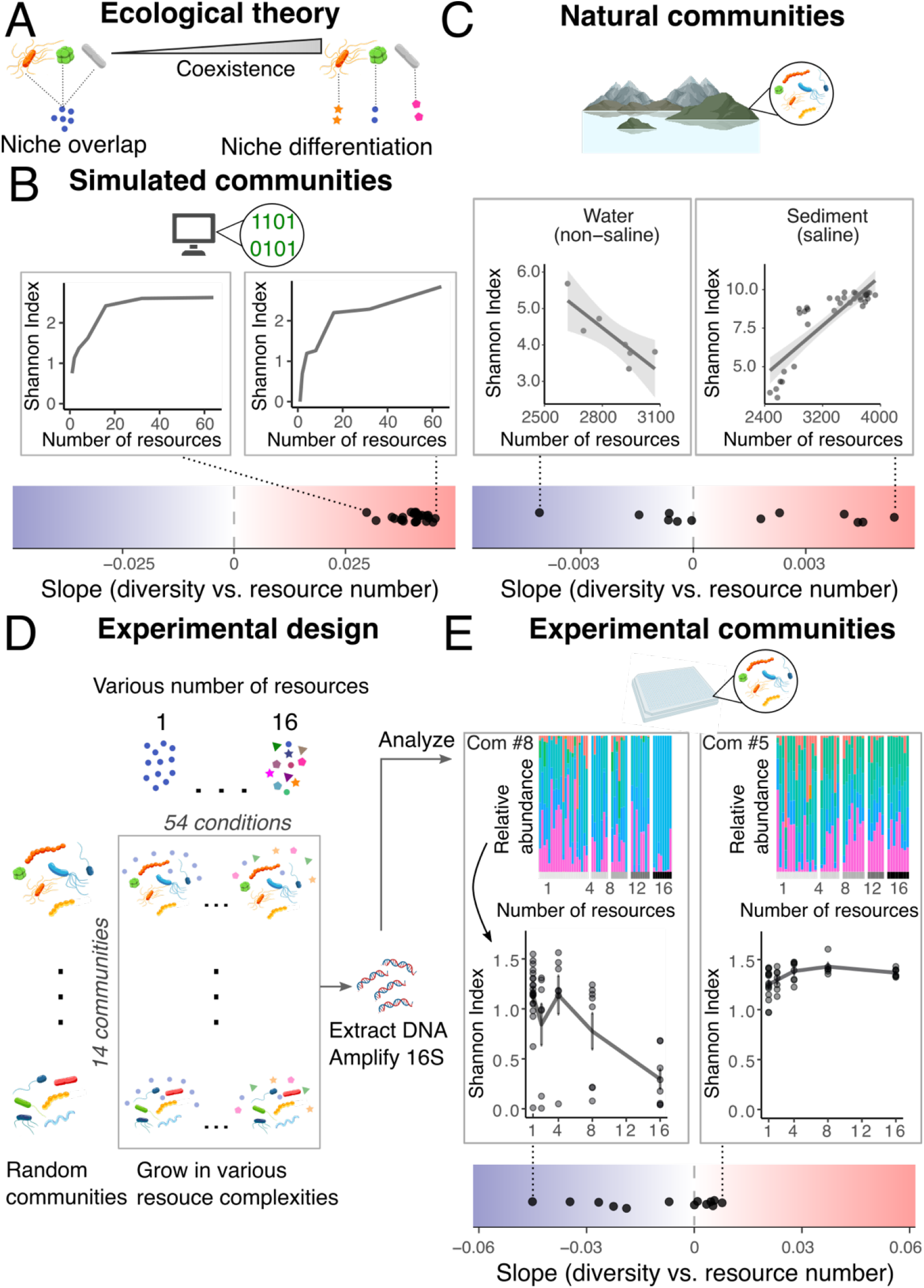
Microbial biodiversity often declines with increasing resource diversity across natural and experimental communities. **A**. When microbial species prefer different resources, higher resource diversity is expected to promote coexistence by reducing competition. **B**. This expectation is supported by simulations using a Consumer–Resource Model (**Supplementary Text 1**; 25 simulated communities). (**Top**) Example communities showing the weakest and strongest biodiversity responses to increasing resource number. (**Bottom**) For each community, biodiversity response was quantified as the slope (β_1_) of a linear model (Shannon index = β_0_ + β_1_ × number of resources + ε). The mean slope across all simulated communities is shown in **Supplementary Fig. 1A** (Shannon index) and **Supplementary Fig. 1B** (richness). **C**. In natural microbial communities, biodiversity does not consistently increase with resource diversity. Data from the Earth Microbiome Project^15^ (462 samples across 13 ecosystems) combine 16S-based diversity estimates with mass spectrometry-derived resource profiles (Methods). (**Top**) Example ecosystems showing positive and negative relationships between resource diversity and microbial biodiversity (full results in Supplementary Fig. 1C). A similar pattern is observed using richness instead of Shannon diversity (Supplementary Fig. 1D). Notably, the most data-supported—’animal distal gut (non-saline)’—shows a significant negative correlation. (**Bottom**) Extending this analysis to all 13 ecosystems, we derive the slope (β_1_) of Shannon diversity index vs. resource number as described in Fig. 1B. **D**. To experimentally test the effect of resource complexity on microbial diversity, we used 14 bacterial strains (Supplementary Data 2) to assemble 14 random synthetic communities—seven with 6 members and seven with 8 members. Each community was cultured across five levels of resource diversity (1, 2, 4, 8, or 16 carbon sources), totaling 54 conditions per community. **E**. In our controlled experiments, microbial biodiversity generally decreased with increasing resource diversity, while the magnitude of this effect varied across communities. (**Top**) Communities with the strongest biodiversity decrease and increase in response to resource diversity are shown, along with relative abundances of their member strains. Shading indicates mean ± SEM. Results using richness are shown in Supplementary Fig. 1F. (**Bottom**) Extending this analysis to all 14 communities, we derive the slope (β_1_) of Shannon diversity index vs. resource number as described in Fig 1B.

This principle is formalized in the Consumer–Resource Model^7^, which tracks microbial growth and resource dynamics. When species are assigned random metabolic traits and uptake efficiencies, the model predicts that increasing resource diversity allows more species to persist—as naively expected (**Fig. 1B; Supplementary Fig. 1A-B**). Although additional mechanisms such as stochastic processes^8^, viral infections^9^, metabolic interdependencies^10^, and environmental fluctuations^11,12^ have also been proposed to sustain microbial biodiversity, the concept that resource diversity drives species biodiversity remains foundational. Yet despite decades of theoretical support, empirical evidence for this link remains scarce^13,14^, raising the question of whether this long-standing theory truly holds in microbial ecosystems?

As a first step toward addressing this question, we re-analyzed multi-omics data from the Earth Microbiome Project^15^ and found that resource diversity does not consistently correlate with biodiversity (**Fig. 1C**; **Supplementary Fig. 1C-D**): in some ecosystems biodiversity increases with higher resource diversity, while in others, it decreases. However, natural systems are uncontrolled, making it difficult to isolate causal effects. Yet only a few controlled experiments have directly addressed this question^13,14^. Among them, just one—by Pacheco et al.^14^— examined how increasing resource complexity affects diversity in a defined microbial community, finding only a weak biodiversity increase. However, the use of a single community limits the generalizability of these findings.

Here, we systematically tested hundreds of microbial communities and thousands of interactions, and found that—contrary to expectations—microbial biodiversity tends to decline as resource diversity increases.

## Results

### Greater resource diversity often reduces microbial biodiversity

To test the fundamental relationship between resource diversity and microbial biodiversity, we assembled random microbial communities across a range of resource diversities (**Fig. 1D**; see **Methods** for details). Communities were cultivated for nine days under a daily dilution regime to allow them to stabilize (**Supplementary Fig. 2**), and final community composition was assessed using 16S rRNA amplicon sequencing. This yielded 701 sequenced samples across different experimental conditions (**Supplementary Data 1**).

Contrary to predictions from the Consumer–Resource Model, microbial biodiversity declined in a large fraction of communities as resource diversity increased, while increases in biodiversity were rare and typically modest (**Fig. 1E**, compared with Fig. 1B; **Supplementary Fig. 1E–F**; **Supplementary Fig. 3**). These trends mirrored patterns observed in natural ecosystems, though the declines were more consistent and pronounced under controlled conditions (compared with Fig. 1C). Together, these findings challenge the ecological expectation that higher resource diversity promotes coexistence and raise the question of what drives this widespread biodiversity loss.

### Resource diversity fuels competition, which reduces coexistence

Microbial communities are strongly shaped by interactions between their constituent strains. We hypothesized that the observed loss of biodiversity might result from a fundamental shift in these interactions triggered by increasing resource diversity. To test this, we systematically co-incubated all pairwise combinations of the 14 strains used in our synthetic communities under a wide range of resource conditions, grouped into three levels of resource diversity: 1, 8, and 16 resources. For each pairwise co-culture, we measured total biomass (OD600) and relative abundances after five daily dilutions to infer interaction outcomes. We also measured how each strain grew in isolation in each environment, allowing us to disentangle intrinsic growth from interaction effects. We then inferred interaction strengths between all strains by fitting individual and pairwise data, along with community data (**Fig. 1E**), to a generalized Lotka–Volterra (gLV) model using Bayesian regression (**Supplementary Text 2**). This yielded interaction matrices for the three levels of resource diversity (illustrated in **Fig. 2A**; **Supplementary Fig. 4**), which, when used to simulate community assembly, reproduced the observed decline in biodiversity (**Supplementary Fig. 5**), supporting the validity of the inferred interactions. In these matrices, diagonal terms represent self-inhibition (how much a strain limits its own growth; α_self_), and off-diagonal terms capture how strongly a strain inhibits others (α_others_). The α values represent *per-capita* interaction strengths, meaning they are independent of population densities. According to theory, when interspecies inhibition exceeds self-inhibition (α_others_ > α_self_), competitive exclusion will occur, where one strain will outcompete the other^16^. Therefore, if biodiversity loss is driven by pairwise interactions, we would expect increasing resource diversity to shift this balance in favor of exclusion. Interestingly, both self- and interspecies inhibition became weaker as resource diversity increased (**Fig. 2B**, top; **Supplementary Fig. 6, 7**). However, self-inhibition (α_self_) decreased more rapidly than interspecies inhibition (α_others_; **Fig. 2B**, middle), leading to a greater number of predicted competitive exclusions as resource complexity increased (**Fig. 2B**, bottom; **Supplementary Fig. 6C**). Counterintuitively, it is therefore not stronger per-capita inhibition of competitors, but greater individual growth—driven by reduced self-inhibition—that drives competitive exclusion and biodiversity loss. Increased growth leads to larger population sizes, which exert stronger total inhibitory effects on competitors, even as per-capita interspecies inhibition becomes weaker with increasing resource diversity.

**Figure 2:**
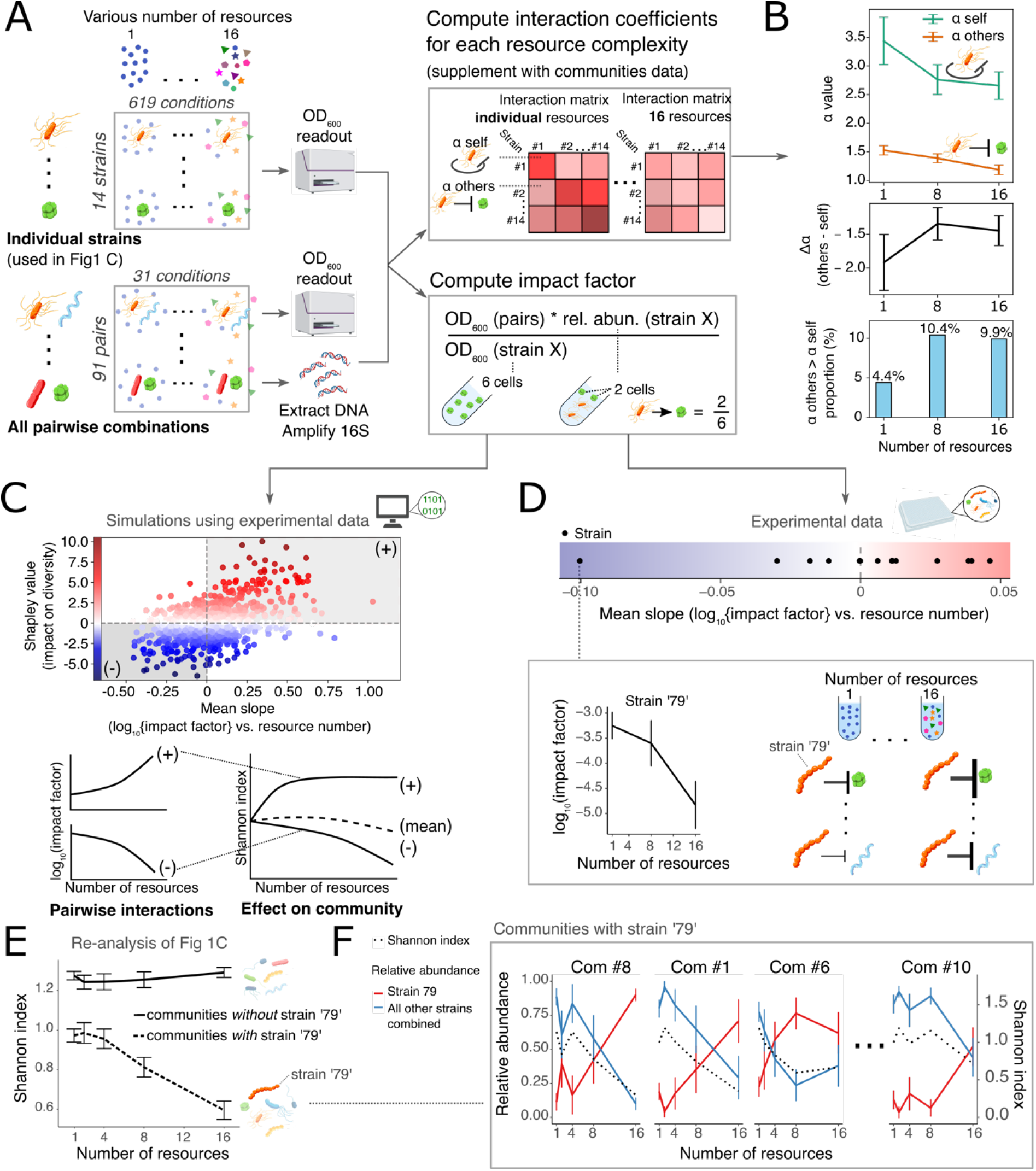
Increasing pairwise competitiveness reduces community biodiversity as resource diversity increases. **A**. Overview of the experimental and modeling workflow. We studied pairwise interactions among 14 bacterial strains across 1, 8, and 16 carbon source conditions. Each pair was tested in an average of 31 conditions, and individual strains were profiled in all their respective conditions, totaling 619 measurements (Supplementary Data 3). OD_600_ and 16S rRNA sequencing were used to quantify growth and composition. Interaction matrices were inferred by applying Bayesian regression to a generalized Lotka–Volterra model (Methods), using observed data to estimate intra-species (α_self_) and inter-species (α_others_) interaction strengths (Supplementary Text 2; results in panel B). Interaction matrices were inferred by a Bayesian regression on a generalized Lotka–Volterra model (Supplementary Fig. 4), allowing comparison of intra-species (α_self_) and inter-species (α_others_) interactions across resource levels (panel B). To assess strain-level contributions to diversity loss, we defined an impact factor based on the change in a strain’s partner’s abundance in co-culture versus monoculture. As exampled, a drop from 6 to 2 yields an impact factor of 0.33. **B**. Resource complexity shifted communities toward competitive exclusion. On average, α_self_ decreased more than α_others_, reducing coexistence potential (top and middle panels). The proportion of interactions where α_others_ > α_self_ increased with resource number (bottom panel). Trend lines show mean ± SEM. **C**. Simulations using noise-perturbed experimental data showed that strains with more negative impact factor slopes (1 vs. 16 resources) contributed more to biodiversity loss. The cartoon illustrates how increased pairwise suppression by a single strain can scale up to reduce overall community diversity. **D**. Strain ‘79’ showed the steepest increase in suppressiveness among experimentally tested strains. For each strain, we fitted log_10_(impact factor) = β_0_ + β_1_ × number of resources + ε across 1-, 8-, and 16-resource conditions (3,398 values total). Strain ‘79’ showed the most negative β_1_. The cartoon illustrates how its suppressive effect increased consistently across interaction partners with resource diversity. **E**. Communities containing strain ‘79’ (n = 8) showed a sharp biodiversity decline with increasing resource diversity, while those without it (n = 6) slightly increased. Trend lines show the mean ± SEM. **F**. In these same communities, strain ‘79’ increased in relative abundance while other strains declined, along with a drop in Shannon diversity. Trend lines show the mean ± SEM. Shown are four representative examples (full data in Supplementary Fig. 12).

The above findings explain the average biodiversity loss with increasing resource diversity, but the strength of this effect varied across communities (Fig. 1E). This suggests that community composition influenced the outcome, with some strains playing a larger role than others. To identify which strains drive biodiversity loss, we defined an ‘impact factor’—a summary metric combining all four interaction coefficients from a pairwise interaction (2×α_self_ + 2×α_others_) into a single value (**Fig. 2A**; **Methods**). This metric reflects how strongly one strain affects the growth of its interaction partner: a value of 1 indicates neutrality, values below 1 indicate suppression, and values above 1 indicate facilitation.

We hypothesized that strains whose impact factor decreased—i.e., became more suppressive—as resource diversity increased would be key drivers of biodiversity loss. To test this, we extensively simulated communities using synthetic strain sets generated by adding random noise to the experimentally derived interaction matrices (**Methods**). This approach allowed us to generalize beyond the limited statistical power of the experimental dataset. For each simulated community, we calculated biodiversity under low and high resource diversity and matched this to strain-specific changes in impact factor. Strains that became more suppressive with increasing resource diversity consistently drove greater biodiversity loss, validating our hypothesis (**Fig. 2C**; **Supplementary Fig. 9**).

To confirm these predictions experimentally, we examined our experimental dataset and found that strains becoming more suppressive with increasing resource diversity contributed more strongly to biodiversity loss (**Supplementary Fig. 10**). One strain in particular, ‘79’, stood out—showing a pronounced decrease in impact factor (i.e., increased suppression) as resource diversity rose (**Fig. 2D**; **Supplementary Fig. 11**). Comparing communities with and without strain ‘79’ revealed a striking pattern: those containing it exhibited sharp biodiversity declines with increasing resource diversity, while those without it showed slight biodiversity gains (**Fig. 2E**). In communities with strain ‘79’, its relative abundance increased steadily with resource diversity, eventually dominating at 16 resources as the total abundance of other strains declined (**Fig. 2F**; **Supplementary Fig. 12**). Shannon diversity mirrored this trend, decreasing in parallel with the collapse of the remaining community members. Taken together, these findings show that increasing resource diversity can amplify the suppressive potential of specific strains, which in turn drives biodiversity loss.

This raises two key questions: how common is it for strains to become more competitive as resource diversity increases, and what physiological mechanisms underlie this shift?

### Increased microbial competitiveness with resource diversity is widespread

To assess how often strains become more competitive with increasing resource diversity, we measured our established impact factor metric (illustrated in **Fig. 2A**) for 298 strains against a sentinel strain across a wide range of resource conditions, generating ∼32,000 data points (**Fig. 3A**; **Methods**). As resource complexity increased, most strains became more suppressive (**Fig. 3B**, top; **Supplementary Fig. 13**). However, within this trend, we identified two distinct response types: mildly influenced strains (189) and highly influenced strains (109), the latter showing a much sharper gain in suppression under complex conditions (**Fig. 3B**, bottom; **Supplementary Figs. 14, 15**). Both types were phylogenetically conserved, suggesting that evolutionary constraints shape competitive strategies (**Fig. 3B**, top). Notably, these response types were already distinguishable based on how strains behaved in individual-resource environments (**Supplementary Figs. 16–17**). The responses of highly and mildly influenced strains were strongly quantitative, scaling with the number of available resources. Highly influenced strains started out less suppressive than mildly influenced strains under simple conditions, but became more suppressive as resource complexity increased (**Fig. 3C**). Strikingly, these strains also lost the facilitative effects that were common in individual carbon sources (**Fig. 3B**, top; **Supplementary Fig. 15**).

**Figure 3:**
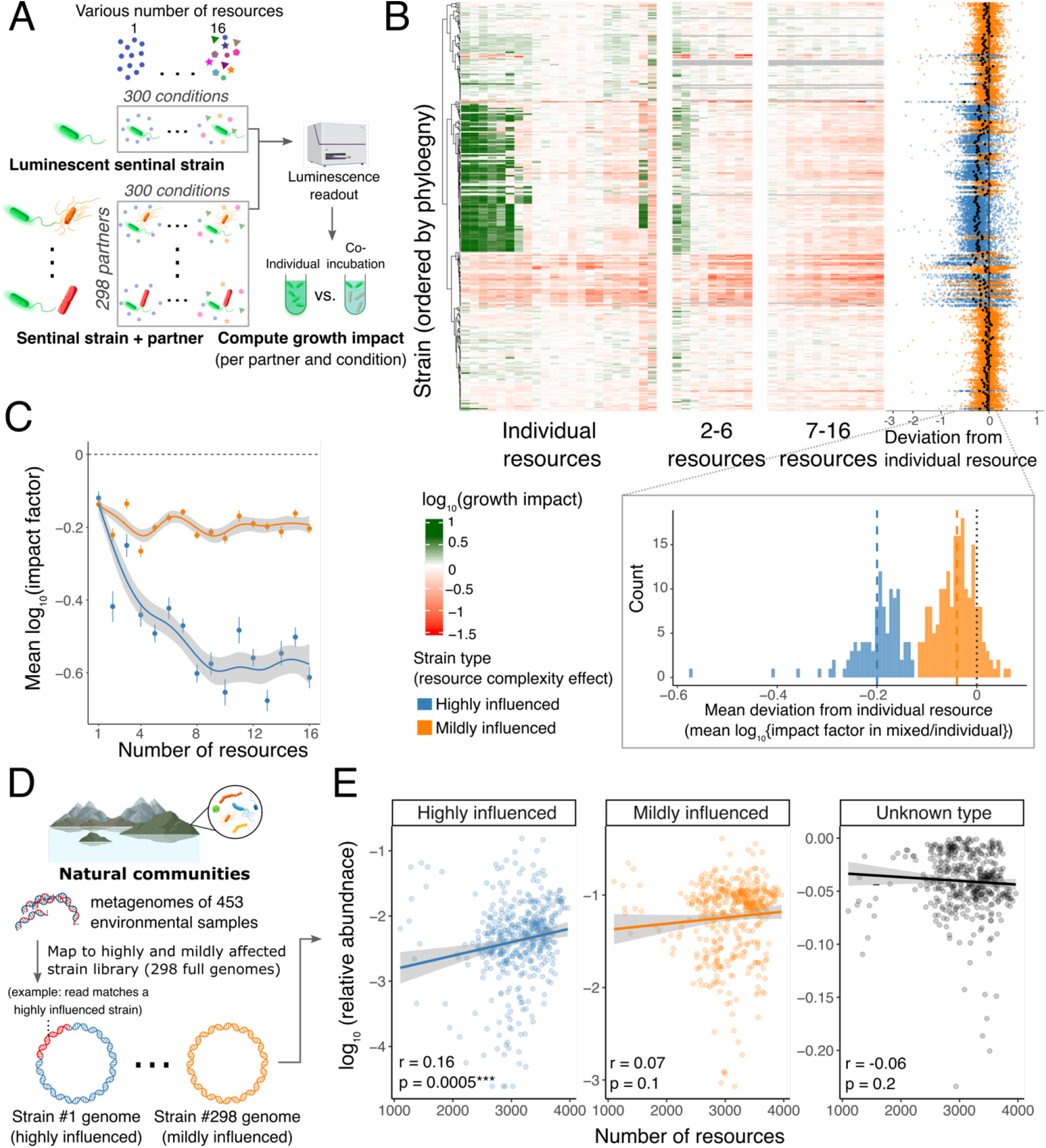
Resource complexity consistently increases microbial competitiveness, particularly among a subset of strains. **A**. We performed a high-throughput experiment using a luminescent sentinel strain (*Pseudomonas aeruginosa PA01*) to quantify microbial interactions. Its growth was measured alone and in co-culture with each of 298 partner strains across 16 levels of resource diversity (1–16 resources), including 22 individual resources and 278 random mixtures (Supplementary Data 4). This yielded 32,216 impact factor measurements quantifying each partner’s effect across conditions. **B**. K-means clustering (Supplementary Fig. 14) classified strains into two types based on their impact factor response: highly influenced (109 strains) and mildly influenced (189 strains). Both types became more suppressive on average with increased carbon complexity, but highly influenced strains showed a much stronger shift. (**Top**) Heatmap shows log_10_(impact factor) across individual resources and selected mixtures (gray = missing data). Strains are ordered by core-genome phylogeny. Right: each strain’s deviation in resource mixtures (≥2 resources) from its mean impact across individual resources. black points represent each strain’s average deviation. In total, 6,556 impact factor values were calculated for individual resources (all shown in the heatmap) and 26,660 for resource mixtures (heatmap and plot on the right). (**Bottom**) Histogram shows the distribution of mean log_10_(impact factor) differences between individual and mixed resources (2–16 resources) for highly and mildly influenced strains. The bimodal pattern reflects the distinct groupings, with dashed lines indicating each group’s mean. **C**. Highly influenced strains show a sharp decline in impact factor, indicating greater suppression, with increasing resource complexity; mildly influenced strains shift only slightly. Lines show mean ± SEM, with smoothed generalized additive model fit and 95% confidence intervals. **D**. Metagenomic reads from 462 Earth Microbiome Project samples across 18 ecosystems were mapped to the full genomes of the 298 tested strains, grouped by their response to resource complexity (highly or mildly influenced; see panel B and Methods). Read counts per group were calculated and converted to relative abundances. Illustration: a metagenomic read mapping to a highly influenced strain. **E**. The relative abundance of highly influenced strains increases with resource complexity across all samples, in contrast to mildly influenced or unmapped reads. Pearson correlation (r) and p-values are shown.

Given the increasing dominance of more competitive strains in our experiments (**Fig. 2F**), we hypothesized that highly influenced strains would also become more abundant in natural ecosystems as resource diversity increased. To test this and extend our findings beyond the lab, we mapped metagenomic data from environmental samples in the Earth Microbiome Project^15^ (**Fig. 1B**) to the genomes of the 298 strains tested in the sentinel assay (**Fig. 3D**). Because highly influenced strains show a phylogenetic signal (**Fig. 3B**), their close relatives are expected to share similar traits. Indeed, only reads mapping to highly influenced strains increased significantly with resource diversity, unlike mildly influenced strains or unmapped reads (**Fig. 3E**). These findings suggest that highly influenced strains not only become more competitive in the lab, but also tend to dominate natural ecosystems as resource complexity increases—reinforcing the ecological relevance of our experimental results.

### Emergent resources uptake raises microbial competitiveness

Microbes often interact by chemically modifying their environment—for instance, by depleting shared resources. Therefore, we examined whether microbial alterations to the growth medium—capturing their metabolic footprint—could reveal the mechanism underlying the observed increase in competitiveness with resource diversity. Specifically, we grew a subset of highly and mildly influenced strains in individual- and mixed-resource settings, filtered the cultures, and collected supernatants. We then compared true mixed-resource supernatants to their equivalent mixtures of single-resource supernatants (illustrated in **Fig. 4A**). If metabolism were unaffected by resource number, both would show similar chemical profiles. However, gas chromatography–mass spectrometry (GC–MS) showed that residual key nutrients were lower in true mixed-resource supernatants, indicating increased carbon uptake with greater resource diversity (**Fig. 4C**; **Supplementary Fig. 18**)—also providing a probable explanation for the stronger microbial growth observed in mixed resources (**Fig. 2B**). This effect was particularly pronounced in highly influenced strains and evident across both consumed and secreted metabolites (**Supplementary Fig. 19**).

**Figure 4:**
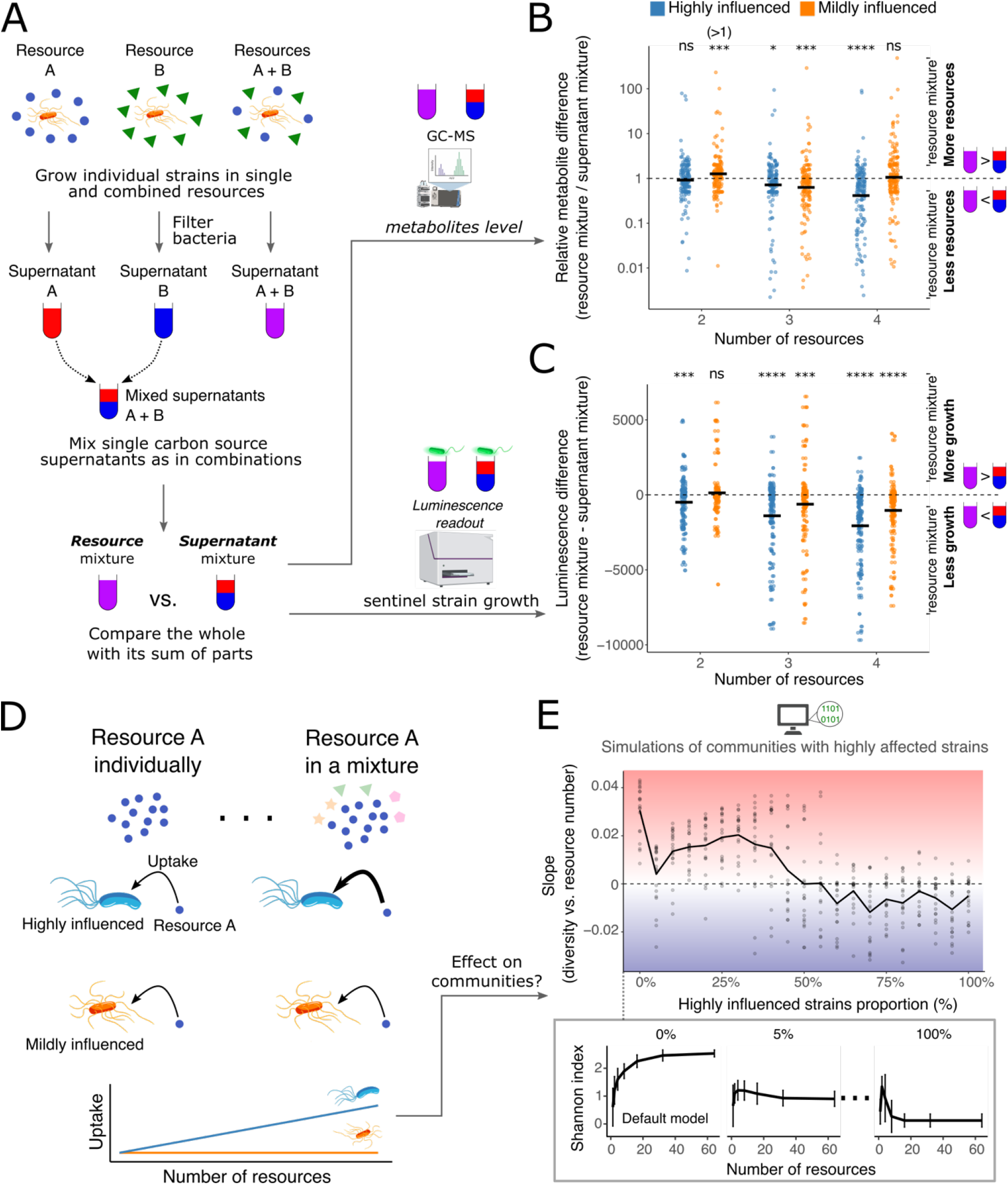
Emergent resource uptake with increasing resource complexity enhances microbial competitiveness. **A**. We tested whether highly influenced strains increase their resource uptake in mixtures beyond what is expected from independent uptake. Strains were grown in individual resources and in mixtures. Supernatants were then extracted, and individual supernatants were combined to create supernatant mixtures (see main text and Methods for details). Metabolic depletion was analyzed by GC-MS, and interaction effects were assessed via sentinel strain luminescence after 24 hours. **B**. Highly influenced strains show greater metabolite depletion in resource mixtures than in their corresponding supernatant mixtures, and this effect intensifies with increasing resource complexity. Shown is the relative change in metabolite peak area (normalized to an internal standard) for 96 identified metabolites, comparing each resource mixture to its matched supernatant mixture. Mann-Whitney U tests assessed whether the median relative change deviated from 1. For mildly influenced strains in two-resource conditions, values >1 indicate less depletion (i.e., lower uptake) in the resource mixture. Significance: ns (p > 0.05), * (p ≤ 0.05), ** (p ≤ 0.01), *** (p ≤ 0.001), **** (p ≤ 0.0001). **C**. Highly influenced strains suppress the sentinel strain more strongly in resource mixtures than in supernatant mixtures, and this effect increases with resource complexity. Mildly influenced strains show the same trend, but less pronounced. Shown is the change in sentinel strain luminescence after 24 h. Significance levels as in panel B. **D**. Our results suggest that highly influenced strains increase carbon uptake as resource diversity increases. For example, when a focal resource (A) is supplied alone, uptake is similar between strain types. But in a mixture, only highly influenced strains show increased uptake of A. **E**. Highly influenced strain behavior reduces microbial diversity as resource complexity increases in a modified Consumer-Resource Model. We introduced a strain-specific parameter γ that increases uptake affinity (i.e., reduces K_m_) as resource diversity increases (equations in Supplementary Text 3). We simulated communities of randomly generated strains across resource complexities using 21 highly influenced strain fractions (0–100% in 5% steps; 20 replicates each). Shown is the slope (β_1_) from a linear model of Shannon diversity vs. resource number, plotted against the fraction of highly influenced strains. The 0% case corresponds to the default Consumer-Resource Model (Fig. 1A). The trend line shows the mean, ±SD. Results are robust for γ > 0.03 (γ = 0.05 shown; Supplementary Fig. 23), where γ controls increased uptake in highly influenced strains (Supplementary Text 3).

To assess whether this enhanced uptake leads to stronger competition, we compared sentinel strain growth in true mixed-resource supernatants to its growth in corresponding mixtures of single-resource supernatants. In line with the GC–MS results, the sentinel strain grew markedly less in true mixed-resource supernatants, especially those from highly influenced strains and under higher resource diversity (**Fig. 4C**). This reduction was not due to toxicity, as nutrient replenishment restored growth in nearly all cases (**Supplementary Fig. 20**). Together, these results show that highly influenced strains increase resource uptake as resource diversity rises, leaving fewer nutrients available and creating a more competitive environment for other microbes (illustrated in **Fig. 4D**).

To test whether this mechanism is sufficient to explain the observed biodiversity loss (**Fig. 1D**), we incorporated increased resource uptake of highly influenced strains into the classical Consumer–Resource Model (**Supplementary Text 3**; **Methods**). Across simulations varying resource complexity and the proportion of highly influenced strains, the modified model frequently predicted a decline in biodiversity with increasing resource complexity—unlike the original model—with stronger effects as the fraction of highly influenced strains increased (**Fig. 4E**; **Supplementary Figs. 21– 22**). This trend held across a wide range of uptake increases (**Supplementary Fig. 23**). Notably, even a small proportion of highly influenced strains was enough to limit the expected biodiversity increase with resource diversity (compared to communities without them), resulting in the dominance of these strains (**Supplementary Fig. 22**). This outcome closely aligns with our experimental results, where the decrease in diversity under high-resource conditions was largely driven by a single strain (‘79’; **Fig. 2A–C**).

Collectively, our results show that a distinct subset of strains with synergistic carbon uptake becomes increasingly competitive as resource complexity rises—ultimately reducing microbial diversity. An emergent physiological trait at the individual level thus drives the unexpected biodiversity loss at the community scale.

## Discussion

Our findings show that, contrary to ecological intuition and widely used models, increasing resource diversity in microbial communities often reduces biodiversity. Since biodiversity is closely linked to ecosystem resilience^17^, this result has important implications. One particularly striking example is in the context of dietary guidelines, which often associate food variety with greater gut microbial richness^18^—yet our results suggest that this strategy may backfire. More broadly, biodiversity loss due to nutrient enrichment (e.g., fertilization) is frequently observed in forest^19,20^, agricultural^21^, and aquatic ecosystems^22,23^, though these cases typically involve increases in nutrient quantity rather than diversity. These outcomes are typically explained by limiting-factor trade-offs, such as shifts in competition toward light^23,24^. Our study isolated resource diversity alone, keeping total nutrient concentration constant, and still observed systematic diversity loss. This demonstrates that resource complexity itself can be sufficient to drive competitive exclusion.

Conventional ecological models often ignore microbial physiology^4,25^. We show that accounting for it can radically alter predicted outcomes and improve model accuracy, as physiological shifts can cascade to reshape entire ecosystems. This finding is surprising within an ecological framework, yet the underlying physiological pattern— emergent changes in microbial resource uptake—is consistent with mounting evidence: microbes exhibit markedly different behavior in mixed-resource environments, including diauxic shifts and lag-phase dynamics^26^, and distinct, unexpected gene expression profiles^27^. While microbial physiology is undeniably complex, we find that viewing it at a coarse-grained level reveals a surprisingly simple and measurable link to competitive behavior—one that is sufficient to explain community-level patterns. This echoes previous work showing that even rough measures of cellular regulation can strongly predict bacterial growth rates^28^. Together, this argues for tighter integration between microbial physiology and ecological modeling. Doing so offers a more realistic view of how microbial ecosystems behave— and why they sometimes defy expectations.

Finally, our finding reopens a fundamental question: how is microbial coexistence maintained despite competition for the same resources? While recent theories offer new perspectives on this paradox^8–12^, it remains unclear whether existing frameworks —alone or in combination—are sufficient to explain the diversity patterns observed in nature, or whether additional, undiscovered mechanisms shape microbial ecosystems in ways we have yet to uncover.

## Methods

### Media and Bacterial Culture

Bacteria were pre-cultured from frozen stocks in nutrient medium (NM) composed of 1% yeast extract and 1% soytone (both Sigma-Aldrich) for 24 h at 30 °C in shaking (225 rpm). Following pre-culturing, cultures were diluted 1:5 in PBS and inoculated into M9 medium supplemented with the relevant carbon sources (see **Supplementary Table 1** for full list and vendor details) and 1:100 diluted NM. Inoculation was performed using an acoustic liquid handler (Echo 525, Beckman Coulter) by transferring 25 nL of 1:5 diluted culture into 40 µL of media per well. For both community and pairwise experiments (Figs. 1 and 2), cultures were passaged daily by diluting 1:27 into fresh media (1.5 µL into 39 µL). Communities were sampled on days 9 and 10 (after 8 and 9 passages, respectively), and pairwise interactions were sampled on day 5 (after 4 passages). All experiments were conducted in 384-well plates. Community and pairwise experiments were carried out in transparent plates with lids (VWR), while sentinel strain experiments were performed in black plates with lids (Sarstedt, Lumox model) optimized for luminescence readouts.

### Resources and their Mixtures

A total of 22 individual resources were used (**Supplementary Table 1**). Resources were dispensed into 384-well plates containing M9 media without carbon sources using an acoustic liquid handler (Echo 525, Beckman) prior to microbial inoculation. For all resource conditions (individual or mixtures), a final concentration of 0.2% (mass/volume) was maintained. Mixture conditions ranged from 2 to 16 resources and were assigned randomly.

### Strain library and whole-genome sequencing

#### Community and pairwise experiments

For all community and pairwise experiments, we used 14 bacterial strains with distinct 16S rRNA gene sequences, enabling unambiguous taxonomic identification (strain names and 16S sequences listed in **Supplementary Data 2**). These strains are part of a *Caenorhabditis elegans*-associated isolate collection of the Ratzke lab and represent a subset of the 16 strains previously used in a similar context^29^.

#### Sentinel strain experiments

For the sentinel strain experiments, we used 298 bacterial strains isolated from the gut of *Caenorhabditis elegans* (**Supplementary Data 5**). Strains were provided by Dr. Christoph Ratzke and Prof. Hinrich Schulenburg.

#### Bacterial whole-genome sequencing

Of the 298 strains used in this study, 92 had been previously sequenced and assembled (**Supplementary Data 5**; courtesy of Prof. Hinrich Schulenburg). The remaining 206 strains were sequenced as part of this work. Genomic DNA was extracted using the peqGOLD Bacteria DNA Kit (VWR). Whole-genome libraries were prepared with the NEBNext Ultra II FS DNA Library Prep Kit for Illumina (New England Biolabs), using dual index oligos for barcoding. To reduce reagent use and cost, the protocol was miniaturized to 1:7 of the standard volume. This scaled-down protocol was benchmarked against the standard 1:1 protocol and produced comparable results. Libraries were sequenced on an Illumina NovaSeq platform to a mean depth of

∼120×. Raw reads were trimmed using fastp (v0.23.2)^30^ and assembled de novo with SPAdes (v3.15.5)^30^ using the --isolate flag and default error correction. Assemblies were generated using 24 threads per sample. Assembly quality was assessed with QUAST (v5.2)^32^, and metrics were summarized across strains. Contamination was evaluated at the contig level using DIAMOND^33^ followed by MEGAN^34^ with GTDB taxonomy. Genomes were excluded if ≥10% of contigs were assigned to taxa inconsistent with the expected lineage, based on contradictions at or above the genus level. A core genome phylogeny of the strain collection was constructed using panX^35^ (default bacterial parameters, 95% core gene threshold). Sequencing statistics in **Supplementary Data 6**.

### Luciferase tagging of the sentinel strain

Pseudomonas aeruginosa PA01 (‘PA01’) was genomically transformed with the luxCDABE operon using the mini-Tn7 insertion system as previously described106. Briefly, the plasmids pUC18T-mini-Tn7T-Gm-lux (Addgene #64953) and pTNS2 (Addgene #64968) were curated from E.coli DH5a using a plasmid extraction kit (peqGOLD Plasmid MiniPrep Kit II; VWR). PA01 cells were made competent for transformation by treatment with a sucrose 300 mM solution PA01, and were then electroporated with 50 ng of the curated pUC18T-mini-Tn7T-Gm-lux and pTNS2 plasmids. After recovery in 1 ml of LB at 30 °C, the electroporated cells were plated on an LB-agar+Gm30 plate. Plates were incubated for 24 hours at 30 °C, and colonies were tested for their bioluminescence activity using a plate reader. Positive colonies were then further grown in a selective medium (LB+Gm30), and were stocked in 25% glycerol at −80 °C.

### Community composition based on 16S rRNA gene sequencing

Amplicon-based 16S rRNA gene sequencing was used to profile bacterial community composition in pairwise and multispecies experiments.

### DNA extraction

Genomic DNA was extracted using a miniaturized protocol in 384-well plates. Pellets were resuspended in TES buffer, and 10 µL was transferred to fresh PCR plates. Lysis was initiated by adding metapolyzyme (in PBS) and ReadyLyse (Lucigen, 1:5 in TES), dispensed via an ECHO acoustic liquid handler, followed by overnight incubation at 30 °C with shaking (1,350 rpm). The next day, Proteinase K (New England Biolabs, 1:2 dilution of 20 mg/mL stock) and 20% SDS (1% final) were added, followed by incubation at 55 °C for 30 min. RNase A (Omega Bio-Tek, 1:10 dilution) was added and incubated for 5 min at room temperature. Proteinase K was inactivated at 95 °C for 5 min. Samples were centrifuged (4,000 rpm, 10 min), and supernatants were collected. Lysates were diluted 1:6 and then 1:3 in nuclease-free water. DNA concentrations were measured using the PicoGreen assay, and 0.1–10 ng was used as template for 16S amplification.

### 16S library preparation and sequencing

Full-length 16S rRNA genes were amplified using a barcoded PCR protocol with primers 27F and 1492R (barcode sequences listed in **Supplementary Data 7**), followed by sequencing on an Oxford Nanopore platform. PCR-grade water and 2× Hot-Start PCR Master Mix (Biotechrabbit) were combined in ECHO-qualified plates, and 4.4 µL of reaction mix was dispensed into 384-well PCR plates using an ECHO acoustic liquid handler (Beckman Coulter). After centrifugation, 50 nL of barcode-specific primers and 500 nL of genomic DNA were added to each well using the same system. Thermocycling was performed as follows: 94 °C for 5 min; 29 cycles of 94 °C for 60 s, 55 °C for 60 s, and 72 °C for 2.5 min; followed by 72 °C for 10 min and a final hold at 4 °C. Barcoded amplicons were pooled per tube, and libraries were prepared using Oxford Nanopore’s short-read amplicon workflow according to the manufacturer’s protocol, with halved reaction and bead-cleanup volumes. Sequencing was performed on a MinION device (Oxford Nanopore Technologies).

### Read mapping and relative abundance estimation

Raw Nanopore reads were split into 100 parts using seqkit^36^ and filtered by quality (Q ≥ 15) and length (1,500–1,700 bp) using NanoFilt (v2.8.0)^37^. Demultiplexing was performed with a custom barcode set using minibar (https://github.com/calacademy-research/minibar), allowing edit distances up to 6 bases. Reverse complementation, where needed, was performed using a custom Python script (rev_cmplt_minibar_output.py). Strain-specific 16S consensus sequences were generated by pooling reads across replicates, filtering (Q ≥ 18), and aligning the top 1,000 reads per strain with MAFFT (v7.508)^38^. Consensus sequences were computed using a custom script applying a 90% agreement threshold and IUPAC ambiguity codes, and then combined into a FASTA file to serve as a custom reference database. Chimeric reads were removed using vsearch (v2.21.1)^39^ with --uchime_ref against this reference. Non-chimeric reads were mapped to the database using minimap2 (v2.26)^40^ with -ax map-ont. Alignment files were sorted and indexed using samtools (v1.17)^41^, and per-sample read counts were generated using samtools idxstats. The final count matrix was compiled with a custom script (make_count_table_updated.py). All custom scripts and workflow code are available at https://github.com/orshalevsk/Emergent_Biodiv_loss.

### OD600 and bioluminescence measurement

#### OD600

A 384-well plate containing bacterial cultures was mixed for 15 s at 2,000 rpm using a Mixmate plate shaker (Eppendorf), then transferred to a FLUOstar Omega plate reader (BMG Labtech) to measure absorbance at 600 nm.

#### Bioluminescence

Bacterial cultures were transferred to a Lumox 384-well plate (Sarstedt) for bioluminescence measurement. First, 2.5 µL of NM solution was dispensed into each well. Then, 20 µL of bacterial culture was reverse-dispensed into the same wells using a VIAFLO 384 liquid handler (Integra) and mixed with NM for 30 s at 1,500 rpm using a Mixmate plate shaker (Eppendorf). The Lumox plate was incubated at 30 °C in a FLUOstar Omega plate reader (BMG Labtech) for 2 min to allow NM to stimulate bacterial metabolism (notably in *Pseudomonas aeruginosa*), enhancing bioluminescence signal consistency. Bioluminescence was then recorded using the plate reader’s luminescence emission filter, with a gain setting of 3,600 and an exposure time of 1 s.

### Experimental design

#### Communities, pairwise and individual strains experiments

Pre-cultures were prepared as described above (Media and Bacterial Culture), then dispensed into 384-well plates containing M9 media with the appropriate carbon sources and 1:100 NM. Metadata for all conditions and strain combinations used in the community experiment are provided in **Supplementary Data 1**; metadata for pairwise and single-strain experiments are provided in **Supplementary Data 3**. All experimental conditions were fully randomized across wells and plates. At the end of each experiment, bacterial density (OD_600_) was measured for all wells, including individual strains, pairwise combinations, and communities. Only pairwise and community samples were further processed for 16S sequencing: samples were centrifuged at 4,000 rpm for 5 min, supernatants were removed, and pellets were stored at –80 °C. DNA extraction and 16S rRNA gene sequencing procedures are described below (DNA extraction and 16S sequencing). Individual strain growth was measured after 72 hours (carrying capacity, OD_600_), pairwise cultures were serially passaged for five days, followed by a final 24-hour incubation before OD_600_ measurement and 16S profiling and communities were serially passaged for 9 days, followed by a final 24-hour incubation before OD_600_ measurement.

#### Sentinel strain experiments

The full experimental setup for sentinel strain co-cultures is described above (see Media and Bacterial Culture and Resources). Briefly, the sentinel strain and 298 co-cultured strains were grown overnight in NM medium and diluted 1:5 in PBS (without washing). Cultures were then dispensed into 384-well plates containing the appropriate media and resource conditions. Six experimental batches were performed: three using individual resources (biological replicates), and three using distinct mixed-resource conditions (no replicates). In the individual-resource experiments, all 298 strains were co-incubated with the sentinel strain across 22 single-resource conditions. The sentinel strain was also grown alone with 16 replicates per condition. In the mixed-resource experiments, one batch included all 298 strains co-incubated with the sentinel strain in all conditions. In the other two batches, 80 strains were randomly selected per condition to maximize the number of unique resource combinations tested. In these batches, the sentinel strain alone was included with 8 replicates per condition. All conditions were fully randomized across wells and plates, with each experiment distributed over 20–24 separate 384-well plates per batch. Plates were incubated at 30 °C for 24 h before luminescence was measured. The average luminescence of the sentinel strain grown alone in each condition was used to calculate the impact factor of co-cultured strains, as shown in Figs. 2A and 3A. Impact factor values for individual resources represent the average across three replicate experiments (‘5’, ‘6’, and ‘7’). Full experimental conditions and metadata are provided in **Supplementary Data 4**.

#### Supernatant experiments

The full setup for supernatant experiments is described above (see Media and Bacterial Culture and Resources). Briefly, selected highly influenced and mildly influenced strains were grown overnight in NM medium and diluted 1:5 in PBS (without washing). Cultures were dispensed into 384-well plates containing M9 medium supplemented with either individual or mixed carbon sources, with each strain inoculated alone into four adjacent wells per condition (e.g., A1, A2, B1, B2) to ensure sufficient volume for supernatant extraction. Plates were incubated at 30 °C for 24 h. To extract supernatants, technical replicates (four wells per condition) were pooled into 96-well plates using a VIAFLO 384 liquid handler (Integra). Each set of four wells (40 µL per well) yielded a combined volume of 160 µL. The pooled cultures were filtered through 0.2 µm filter plates (PALL) into a new sterile 96-well plate, removing cells and leaving only the cell-free supernatant. This design enabled combinatorial mixing of supernatants to reconstruct mixed-carbon environments. Mixing was performed using a VIAFLO 96 liquid handler by combining supernatants from wells on different plates (e.g., A1 from plate 1 with carbon A and A1 from plate 2 with carbon B were combined 1:1 to generate A+B mixtures). For the sentinel growth assay (Fig. 4C), eight strains were randomly selected—four highly influenced and four mildly influenced—were tested across 66 mixed-resource conditions. For the mass spectrometry experiment (Fig. 4B), a subset of four of these strains (two highly influenced, two mildly influenced) was tested across nine mixed-resource conditions. Full strain lists and conditions are provided in **Supplementary Data 8**.

### GC–MS sample preparation and analysis

A total of 72 samples (20 µL each) were thawed on ice and mixed with 50 µL of internal standard solution containing 60 µM 13C_6_-glucose and 60 µM 3-hydroxybenzoic acid (equivalent to 3,000 pmol each). Samples were dried overnight using a vacuum concentrator (Eppendorf Concentrator) to remove water, then placed in a desiccator over phosphorus pentoxide for 1 h. Derivatization was performed following a protocol adapted from Fiehn et al^42^. Briefly, 50 µL of methoxylamine hydrochloride in pyridine (20 mg/mL, freshly prepared) was added to each sample. Samples were incubated in an ultrasonic bath for 10 min, followed by shaking at 30 °C for 90 min at 1,400 rpm. After a brief centrifugation, 70 µL of MSTFA (N-methyl-N-trimethylsilyltrifluoroacetamide; Sigma-Aldrich) was added. Samples were then incubated at 40 °C for 60 min at 1,200 rpm and left at room temperature for an additional 2 h. Final centrifugation was performed at 13,000 rpm for 10 min at 4 °C (Hettich tabletop centrifuge, swing-out rotor), and 60 µL of supernatant was transferred to GC vials for injection. GC-MS analysis was carried out on a Shimadzu TQ 8040 triple quadrupole mass spectrometer coupled to a high-performance liquid chromatography system. Chromatographic separation was achieved on a Restek Rxi-5Sil MS column. Detailed GC and MS parameters are provided in **Supplementary Data 9**. Data acquisition and peak integration were performed using LabSolutions Insight GCMS software, with manual curation of peak boundaries.

### Earth Microbiome Project data handling

#### *Alpha diversity vs. resource complexity* analysis

Data were retrieved from the Earth Microbiome Project (EMP) study by Shaffer et al.^15^ via Qiita (study ID 13114: https://qiita.ucsd.edu/study/description/13114). We used both 16S-based alpha diversity and predicted microbially derived metabolite richness, as previously computed in Shaffer et al.^15^ (Fig. 2C; Supplementary Figs. 1C–D). Alpha diversity was taken from the column alpha_16s_deblur_nosingletons_rar5k_shannon, which reports Shannon diversity after singleton removal and rarefaction to 5,000 sequences per sample. This depth was identified by the authors as optimal for cross-sample comparison. Resource diversity was taken from alpha_lcms_fbmn_microbial_richness, defined as the number of HPLC–MS peaks >0 in each sample following Feature-Based Molecular Networking (FBMN), and interpreted as microbially associated metabolite richness. Samples were filtered to include only those with non-missing alpha diversity values (477 of 618 total). To ensure statistical robustness, we retained only environments (as defined by EMP level “empo_4”) with ≥6 samples, resulting in 462 samples across 13 distinct ecosystems. We used the “empo_4” level of environmental classification, which provides the most specific ecosystem labels available in the EMP dataset and avoids confounding that can arise at broader levels (e.g., “empo_1”), where ecologically distinct environments —such as animal guts and plants—may be grouped together.

#### Mapping metagenomic reads to highly and mildly influenced strains and abundance estimation

To assess the environmental representation of highly and mildly influenced strains, we mapped metagenomic shotgun reads from the Earth Microbiome Project^15^ to our full genome collection of 298 experimentally characterized strains. Genome assemblies were merged into a single reference database, and a minimap2 index (v2.26)^40^ was created. Reads were aligned using minimap2 with the -ax sr --secondary=no parameters to exclude secondary alignments (pipeline: minimap2_pipline_no_multi_match.sh). Alignment files (SAM) were processed using a custom script (make_count_table_updated_with_mistmatch_filter_db.py) to filter for high-quality matches (--min_mapq 50) and compute per-strain read counts, along with unmapped reads (**Supplementary Data 10**). To ensure sufficient coverage and minimize noise, only samples with a total mapped read count ≥10,000 were retained. This filtering yielded 453 metagenomes spanning 18 distinct ecosystems. For each sample, raw counts were normalized to relative abundances. Strains were categorized as highly or mildly influenced based on their experimentally measured impact on biodiversity, and reads were binned accordingly into highly influenced, mildly influenced, or unmapped groups. Relative abundances for each category were then calculated (**Supplementary Data 11**). These values were integrated with matched metabolomic profiles from the Earth Microbiome Project^15^ for downstream ecological analyses. All custom scripts used in this workflow are available at https://github.com/orshalevsk/Emergent_Biodiv_loss.

### Consumer-Resource model

For the classic Consumer-Resource model see **Supplementary Text 1**. For the highly influenced strains modified version see **Supplementary Text 3**. For the highly influenced strains version, we simulated 420 communities of 50 randomly generated strains across seven resource complexities (1–64; **Fig. 4E**; **Supplementary Figs. 21-23**).

### Inferring interaction matrices from experimental data using the generalized Lotka–Volterra model

See **Supplementary Text 2**. Note that error bars for α_self_ in Figure 2B (middle) may be overestimated as the interaction values are not independent (see Supplementary Fig. 6A).

### Estimating single-species Shapley contributions to biodiversity loss

We used the interaction matrices obtained from experimental data for 1 and 16 carbon sources (methodology described in **Supplementary Text 2**; results in **Supplementary Fig. 4**) and added Gaussian noise (mean = 0, variance = 0.7) to each entry to generate a synthetic set of 14 species resembling the 14 experimentally measured strains. We produced in total 800 such sets of 14 species and took from each set randomly six or eight species respectively (400x six species and 400x eight species). We then simulated all possible subcommunities of these 6 or 8 species using the generalized Lotka–Volterra model under both 1- and 16-resource conditions. Since the interaction matrices were obtained from steady state data, we could not estimate the single species’ *per capita* growth rates and set them to be one instead. Since we are only interested in steady state outcomes which are not impacted by the *per capita* growth rates only ensuring that the systems reached steady state was sufficient for further analysis (**Supplementary Fig. 8**). For each simulated subcommunity, we computed the steady-state Shannon diversity index under both conditions and calculated the change in diversity as: ΔShannon = Shannon_16_ − Shannon_1_. We then computed Shapley values, representing the contribution of each strain in a six- or eight species community to the overall change in Shannon diversity between 1 and 16 resources. Next, using the same interaction matrices (**Supplementary Fig. 4**), we calculated each species’ mean impact factor on the other members of its community under both resource conditions—analogous to the analysis shown in Fig. 2D. Finally, we computed the change in mean impact factor with increasing resource diversity and plotted these values against the corresponding Shapley values for each species (**Fig. 2C**).

## Code availability

The code used in this study is available at https://github.com/orshalevsk/Emergent_Biodiv_loss.

## Data availability

All sequencing data will be deposited in the European Nucleotide Archive (ENA) prior to publication and will be made available upon request during peer review.

## Acknowledgements

C.R. received funding from the European Research Council (ERC) under the European Union’s Horizon 2020 research and innovation programme (grant agreement No. 948753), the Deutsche Forschungsgemeinschaft (DFG, German Research Foundation) grants 468972576 and 540605007, and the Cluster of Excellence EXC 2124 “Controlling Microbes to Fight Infections” (CMFI). O.S. received funding from the DFG (grant 516931136). We thank Hinrich Schulenburg for generously providing a C. elegans strain library, and Andreas Peschel for providing the original Pseudomonas aeruginosa PA01 strain. We also thank the entire Ratzke lab for constructive feedback on the project and the manuscript. The bacterial illustrations in the figures were created by Bala Akaba.

## Author Contributions Statement

O.S. Conceived the study, led the conceptualization, performed all experiments, data analysis, and coding, generated all figures except where noted, and was responsible for writing—original draft and review & editing. C.R. Contributed to conceptualization, developed the generalized Lotka–Volterra modeling framework (including simulations and Bayesian inference), and contributed to writing—review & editing. X.Y. Contributed to conceptualization, extended the Consumer–Resource model to include highly influenced strains, performed related simulations, and contributed to writing— review & editing. J.K. and M.S. Conducted the GC–MS measurements and performed metabolite data analysis.

