## Supplement for "Emergent loss of microbial biodiversity with increasing resource diversity"

**Supplementary Text 3: Consumer-resource model with highly influenced strains modification** 31

#### Supplementary Figures

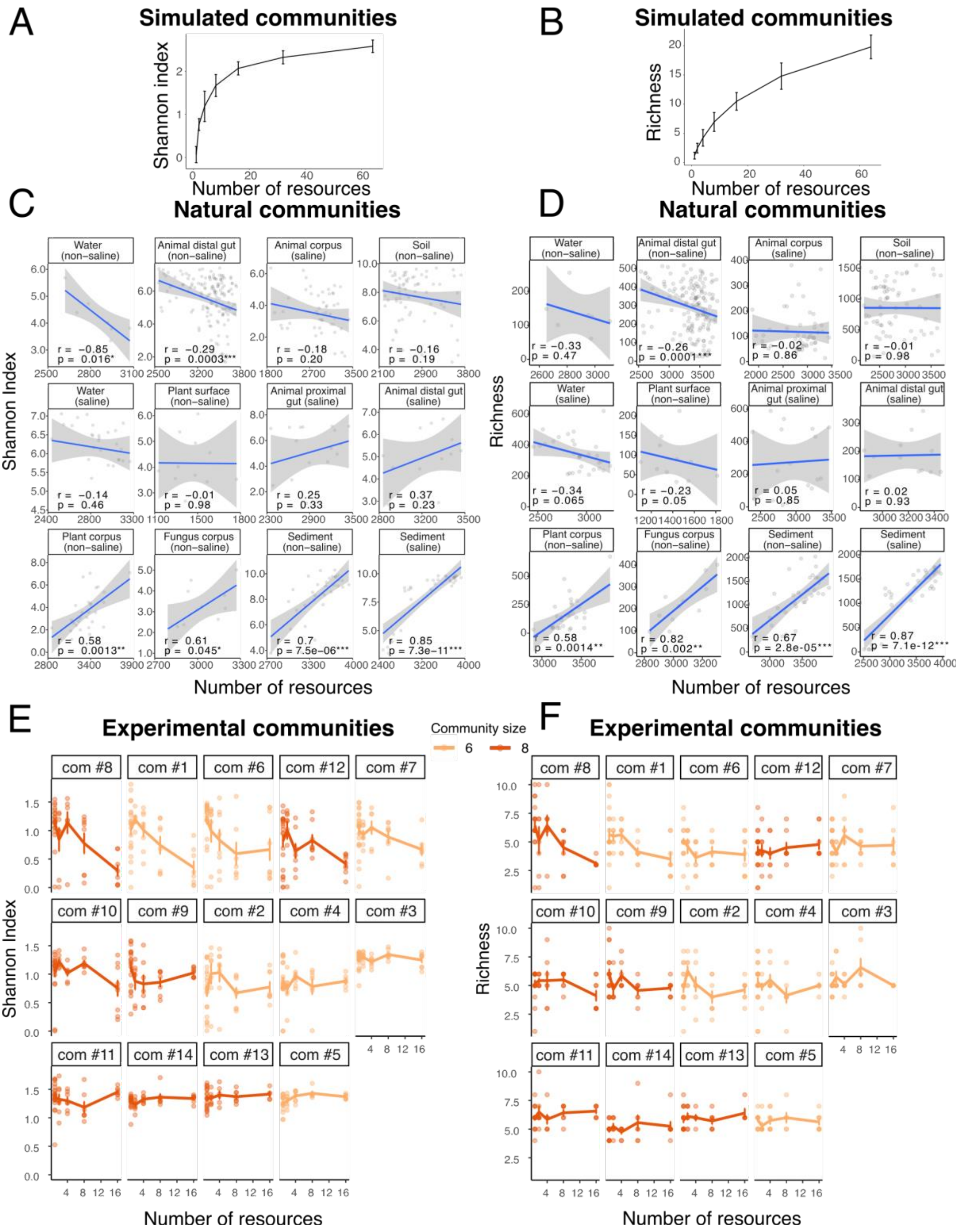

**Supplementary Figure 1: Microbial diversity rarely increases with resource complexity, challenging the intuitive consumer-resource model.** Supplementary results for Fig. 1. **A.** Simulations with the consumer-resource model predict an increase in alpha diversity with niche expansion, shown here using Shannon's diversity index. The trend line represents the average of 25 communities, with error bars indicating ± SD. **B.** Same as panel A but using richness instead of Shannon's diversity index. **C.** In natural communities, the relationship between microbial diversity and resource complexity varies, displaying both positive (consistent with the model) and negative slopes (contrary to it). Data are from the Earth Microbiome Project[1] ('EMP'); 16S metagenomics was used to assess diversity, while mass spectrometry determined the number of resources ('Methods'). Shown are 13 ecosystems at the highest binning level ('empo_4' in EMP nomenclature[1]). The slope of diversity vs. resource number ($\beta_1$) is derived from the linear model: diversity = $\beta_0 + \beta_1 \times$ number of resources + $\varepsilon$. The trend line represents the fitted linear model, with shading indicating the 95% CI. Pearson's correlation analysis (r: correlation coefficient, p: p-value) is shown at the bottom of each plot. Communities are ordered by their slope ($\beta_1$) using Shannon's diversity index (from negative to positive). **D.** Same as panel C but using richness instead of Shannon's diversity index. **E.** In the controlled experiment, microbial diversity responses to resource complexity varied across communities, showing both positive (as predicted by the consumer-resource model) and negative correlations (contrary to it). Data are from 16S sequencing after nine passages (one every 24 hours), followed by 24 h incubation. In total, alpha diversity was calculated for 701 successfully sequenced samples spanning all communities and carbon conditions. The trend line represents the average, with error bars indicating ± SEM. **F.** Same as panel E but using richness instead of Shannon's diversity index.

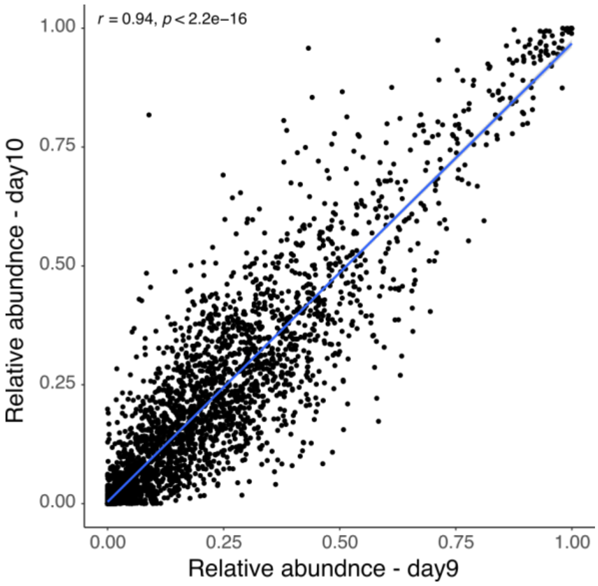

**Supplementary Figure 2: Communities reach a steady state by day 10.** Relative abundances of the same strains within the same sample (strain, community, condition, plate, and well) are shown for day 9 (after eight daily passages into fresh media, followed by 24 h incubation) and day 10 (after nine daily passages and 24 h incubation). Pearson's correlation analysis (r: correlation coefficient, p: p-value) is presented. Data from day 10 were used for all subsequent analyses in this experiment.

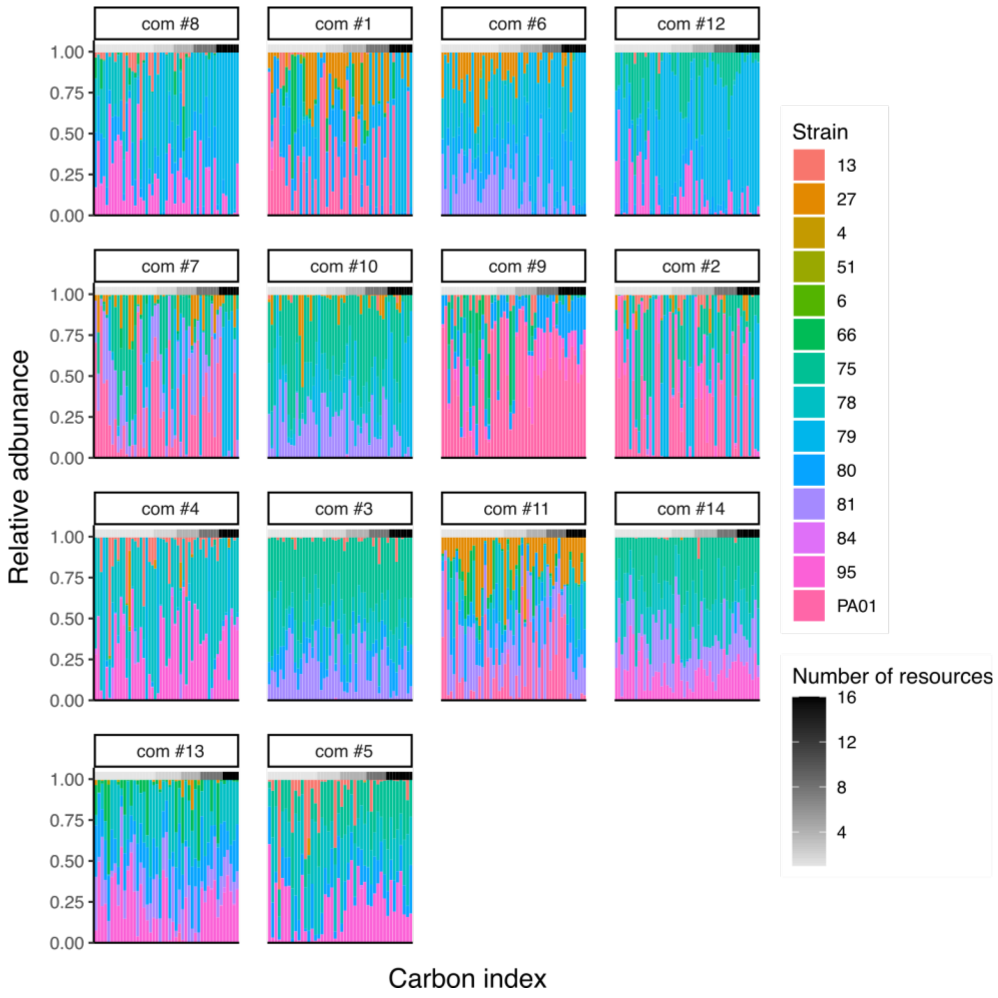

**Supplementary Figure 3: Strain relative abundances across resource complexity in the 14 communities.**

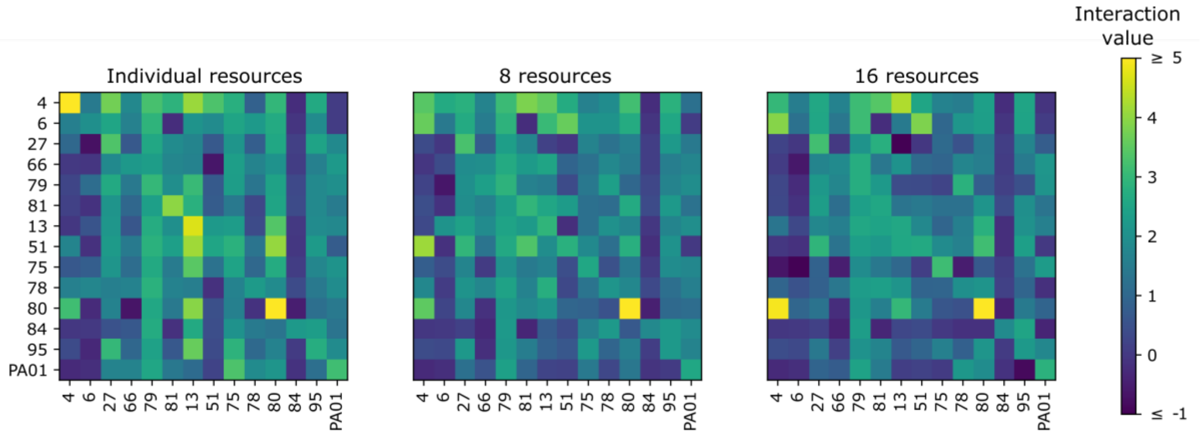

**Supplementary Figure 4: Interaction matrices for 1, 8, and 16 carbon sources, obtained by fitting the experimental data using Bayesian regression (see Supplementary Text 2).** As resource number increases, interaction coefficients decrease for both intra-species (diagonal) and inter-species interactions (off-diagonal).

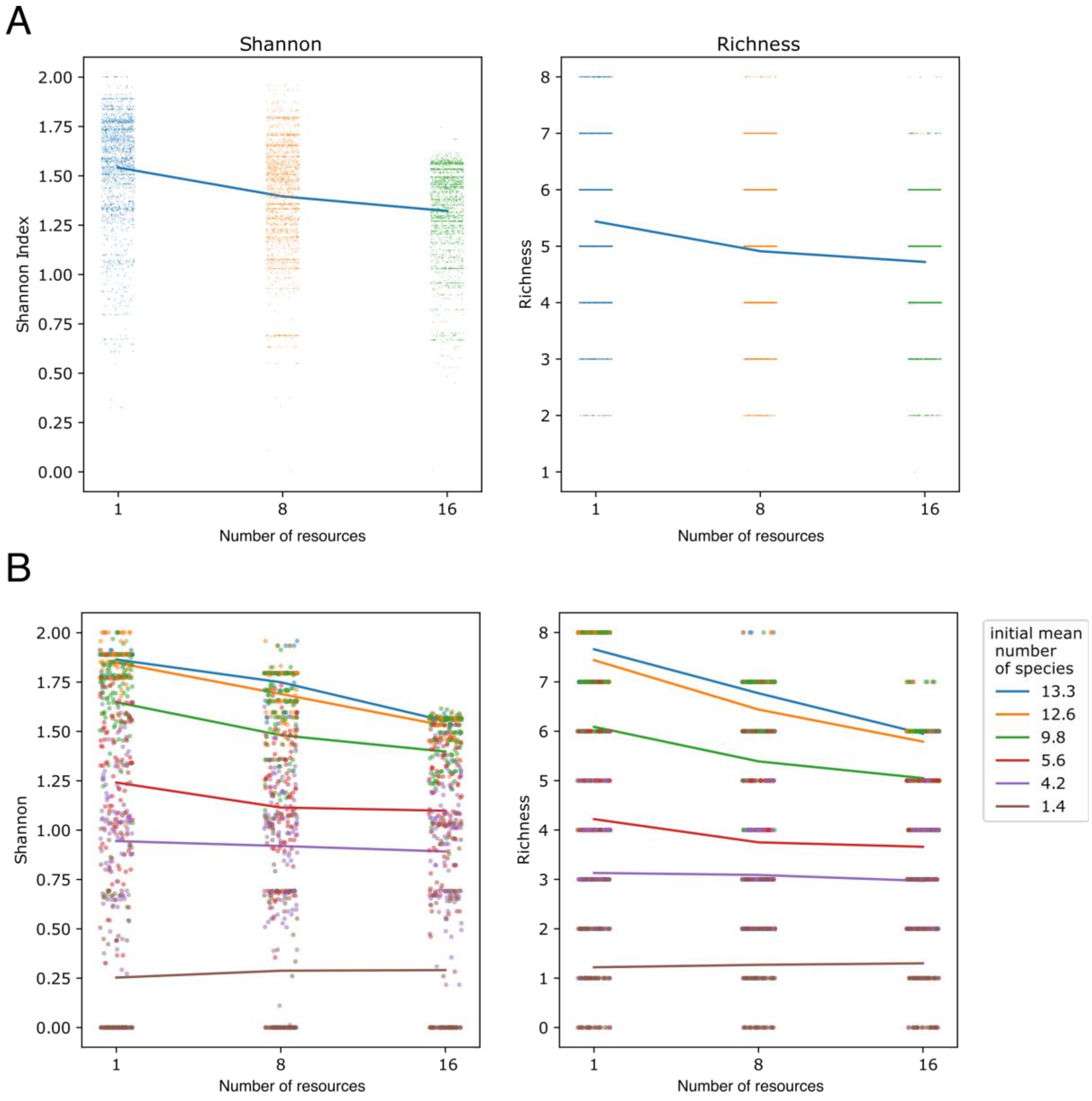

**Supplementary Fig. 5: Simulating microbial communities using interaction matrices derived from experimental data. A.** Simulating 2,000 communities with an average of 8.4 initial species (drawn from the 14 experimental strains) using the interaction matrices shown in Supplementary Fig. 4 reproduces the decline in entropy and richness observed in experimentally measured communities. **B.** Simulations as in panel A but using 100 communities for each fixed initial community size. A reduction in diversity emerges from a certain complexity threshold (approximately 4.2 species) and becomes more pronounced with increasing community size.

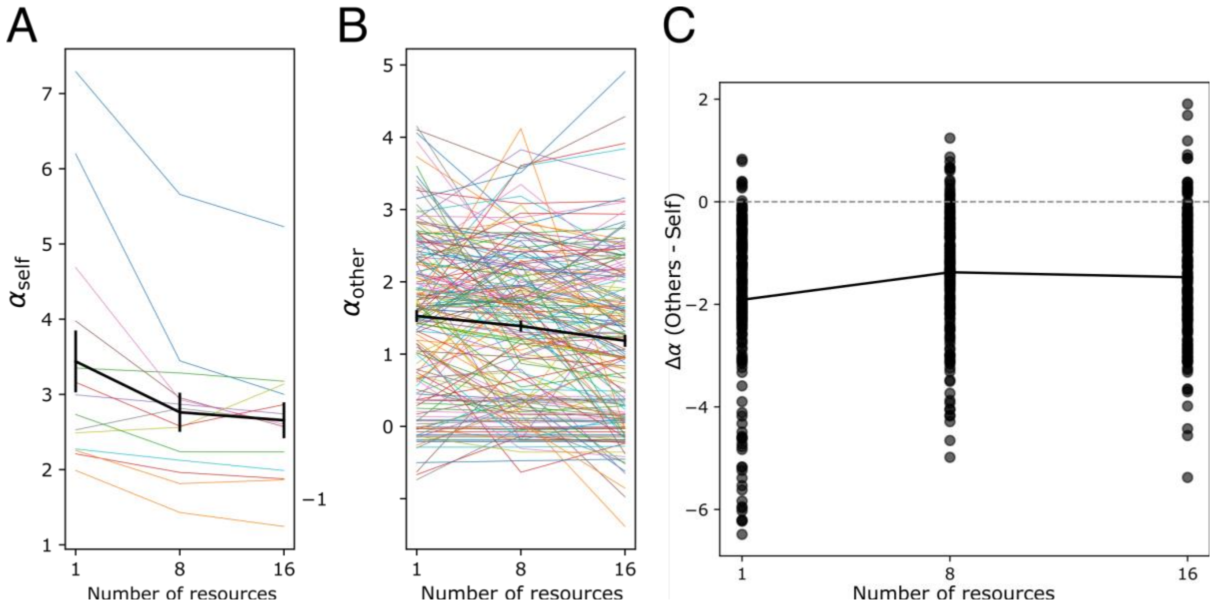

**Supplementary Figure 6: Individual αself and αother values obtained via Bayesian regression, and their difference, plotted as a function of the number of resources. A.** Nearly all $\alpha_{self}$ values decrease with increasing resource number. **B.** $\alpha_{other}$ shows a more complex pattern. Nonetheless, a decreasing trend is also observed on average for $\alpha_{other}$. Black line indicates the mean ± SEM. **C.** Supplement to the bottom panel of Fig. 2B. The system shifts on average toward a competitive exclusion regime at the pairwise level as resource complexity increases. The difference $\alpha_{other} - \alpha_{self}$ decreases with increasing resource number, and the proportion of pairwise interactions where $\alpha_{other} > \alpha_{self}$ (points above the dashed line) increases—conditions under which competitive exclusion occurs.

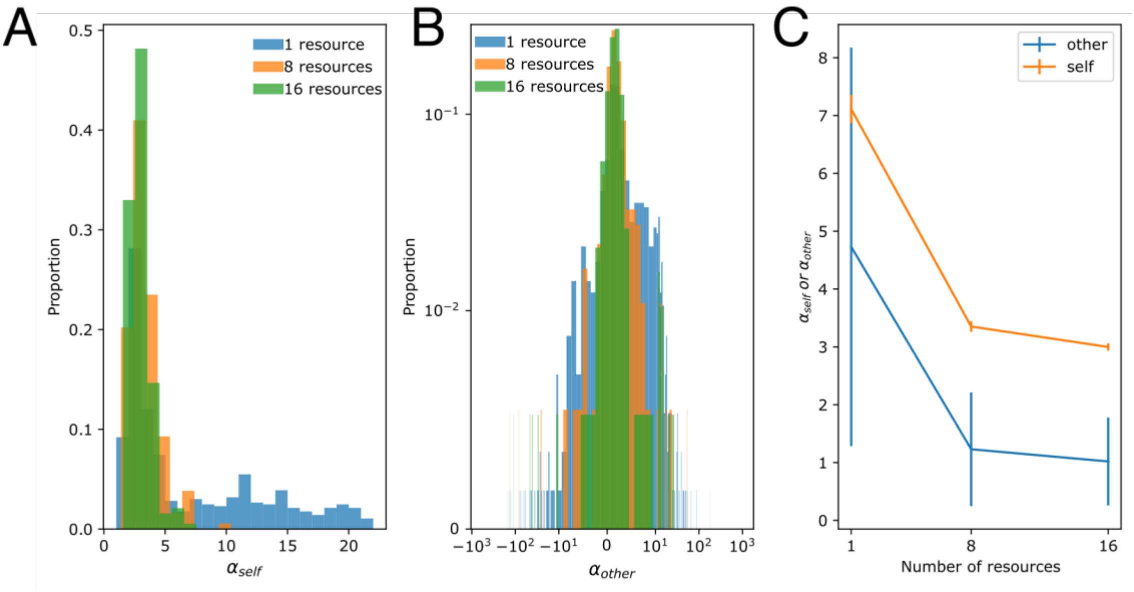

**Supplementary Figure 7: Interaction matrix directly derived from single-species and pairwise growth data.** $\alpha_{self}$ values were estimated as the inverse of carrying capacity, calculated from the final $OD_{600}$ after 72 hours of monoculture growth. $\alpha_{other}$ values were estimated by solving the two-species gLV model under steady-state assumptions: $\alpha_{12} = (1 - \alpha_{11}x_1)/x_2$ and $\alpha_{21} = (1 - \alpha_{22}x_2)/x_1$, using population densities ($x_1$, $x_2$) inferred from each strain's relative abundance multiplied by total community $OD_{600}$ (measured 24 h after five daily passages). To avoid division by near-zero values, strain pairs where either species had a relative abundance ≤ 0.001 were excluded. **A.** Distribution of $\alpha_{self}$ values for 1, 8, and 16 carbon sources. Low population densities in single-resource conditions occasionally produced large, noise-sensitive $\alpha_{self}$ estimates. **B.** This issue was more pronounced for $\alpha_{other}$ values, where small denominators led to extreme values and a broad distribution. **C.** Mean ± SEM of $\alpha_{self}$ and $\alpha_{other}$ across resource richness. The widespread in $\alpha_{other}$ values (panel B) inflated uncertainty in the means. To address this, we adopted a Bayesian regression framework, which allowed us to fit a unified model across all data points while reducing the influence of outliers. Despite the limitations of direct α estimation, the overall trends in panel C are consistent with those obtained using Bayesian regression (Main Fig. 1B).

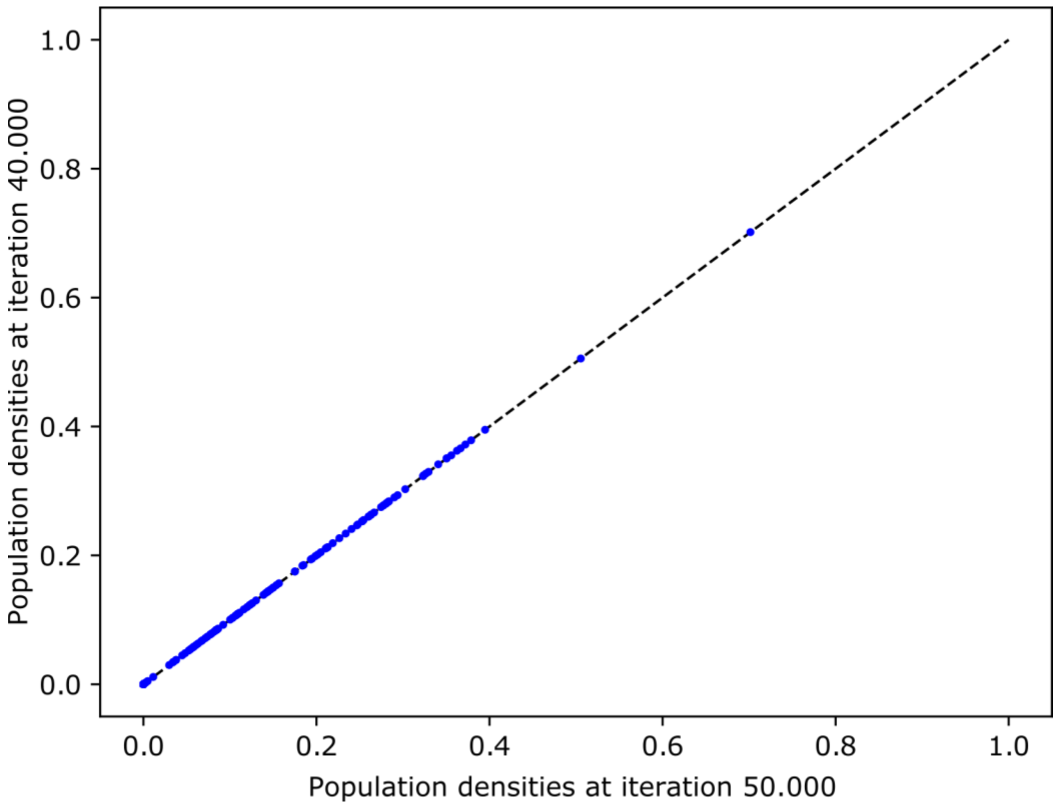

**Supplementary Figure 8: Validation of steady state in gLV simulations.** Related to Methods, chapter '*Estimating single-species Shapley contributions to biodiversity loss'*. Generalized Lotka–Volterra simulations were performed using the same parameters as in Fig. 2C and Supplementary Fig. 5, with interaction matrices from Supplementary Fig. 4. Each simulation run began with six species randomly selected from the full pool of 14. For each matrix, 30 simulations were run for 50,000 iterations. To confirm convergence, species densities at iteration 40,000 were plotted against their densities at iteration 50,000. The overlap between time points indicates that the system had reached steady state. This validation is important because the interaction matrices were derived from steady-state experimental data, but the per capita growth rates (k) could not be inferred and were set to 1. As a result, the model can reliably capture steady-state community structure, but not necessarily population dynamics, which are sensitive to the specific k values.

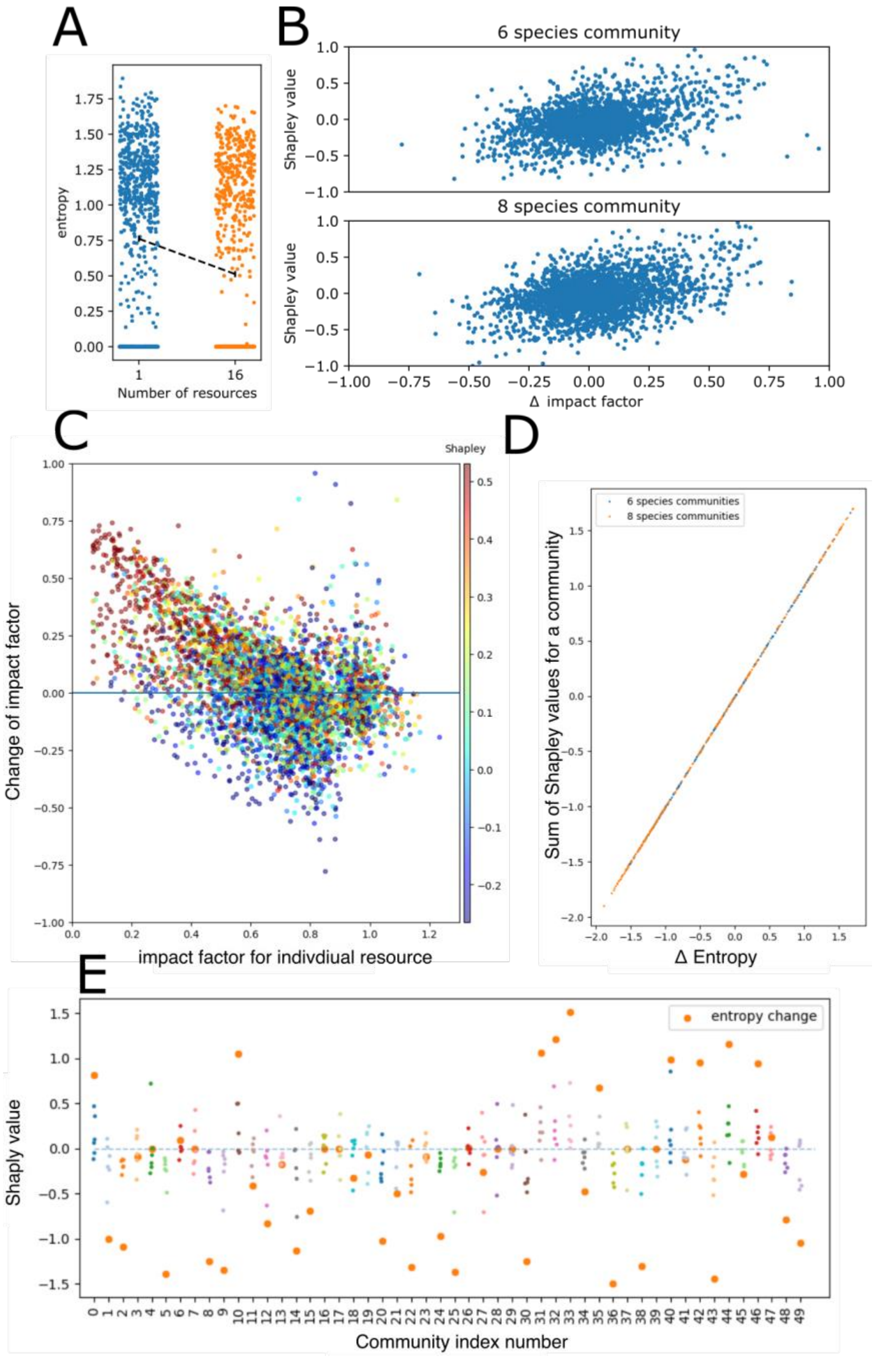

**Supplementary Figure 9: Shapley analysis under varying resource conditions. A.** Effect of resource number on community entropy, interpreted through a Shapley-value lens. Each point represents the entropy of a simulated community under either 1 or 16 resources. The dashed arrow shows the average marginal contribution of increasing resource number to entropy, analogous to a Shapley value. The downward trend reflects a consistent negative effect of resource richness on diversity. **B.** Same analysis as in Main Fig. 2C, shown separately for communities with six and eight species. No significant difference is observed between the two cases. **C.** Both the change in impact factor with increasing resource number and the initial impact factor at one resource influence community diversity. Microbes that are initially weak suppressors but become more suppressive as resource complexity increases tend to reduce community entropy. However, some strains with little or no change in impact factor—and weak initial suppressiveness—still show high Shapley values (blue dots), likely due to interactions with more dominant strains. We speculate that biodiversity loss may result from a combination of increasingly suppressive strains and others that are outcompeted by them. While impact factor is a useful single-strain trait, many strains deviate from this trend, underscoring the emergent and context-dependent nature of community-level properties. **D.** As expected from theory, Shapley values across all species sum to the total change in the selected outcome—in this case, the change in Shannon entropy with increasing resource richness. **E.** Shapley values for the first 50 communities, along with their total change in entropy (sum of individual species' Shapley values; see panel D). In some communities, entropy change is driven by a single species, while in others it results from the combined effects of multiple strains with large absolute Shapley values.

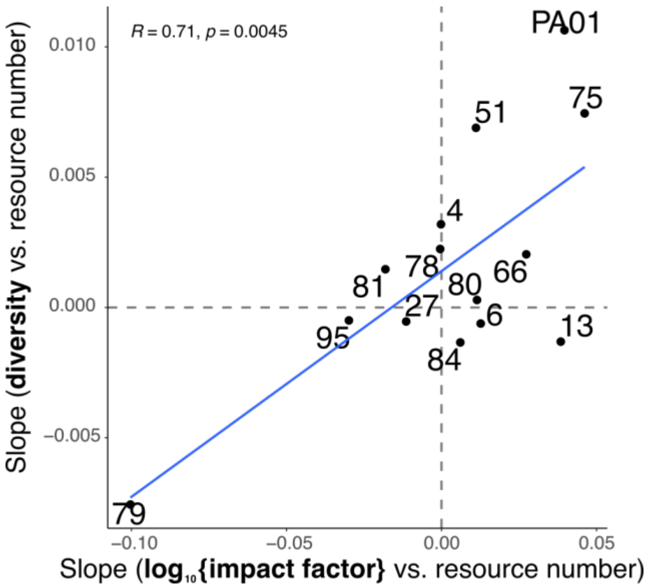

**Supplementary Figure 10: Our experimental results support the same conclusion as the simulations: strains with increasingly negative impact factor slopes drive biodiversity loss, while those with positive slopes have the opposite effect.** X-axis: Slope ($\beta_1$) of $\log_{10}$(impact factor) vs. number of resources, derived from a fitted linear model: $\log_{10}$(impact factor) = $\beta_0$ + $\beta_1$ × number of resources + ε. Y-axis: Slope ($\beta_1$) of Shannon diversity index vs. number of resources, derived from a fitted linear model: diversity = $\beta_0$ + $\beta_1$ × number of resources + ε. The relationship between the two slopes is shown as a fitted linear model. Pearson's correlation coefficient (r) and p-value (p) are indicated in the top left. Strain names are indicated next to each point.

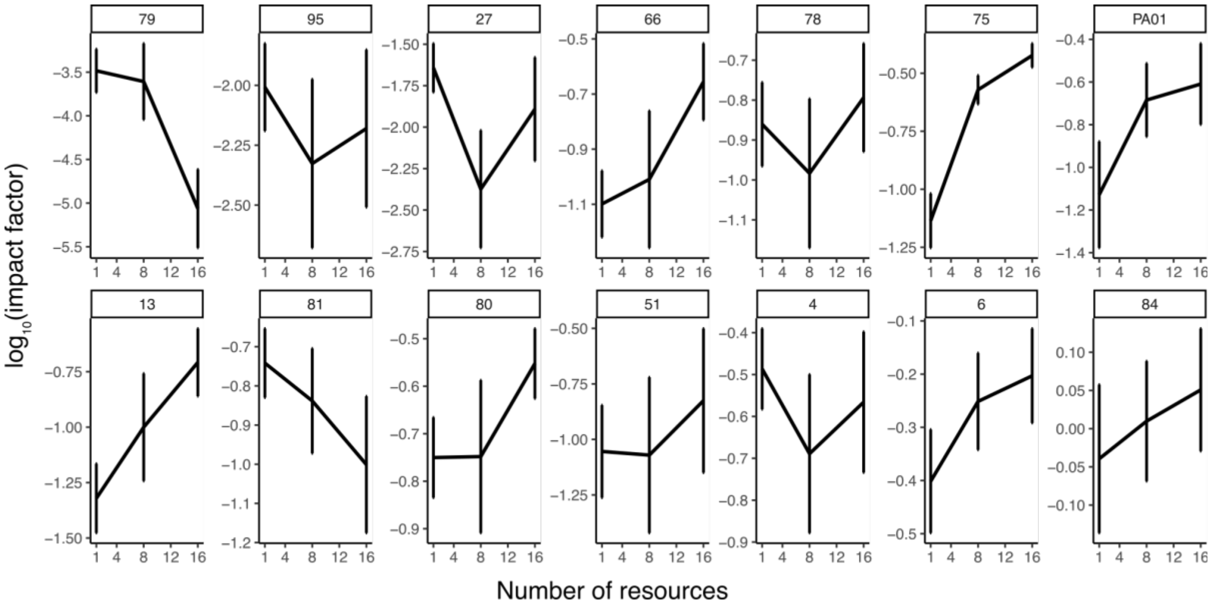

**Supplementary Figure 11: Impact factor of all 14 strains across resource complexity.** Supplementary results for Fig. 2D. Shown is the mean impact factor across 1, 8, and 16 resources for all strains, with error bars indicating ± SEM. Impact factor for each strain was averaged across its interactions with all other 13 strains (i.e., all pairwise interactions) and its growth in isolation (formula and illustration in Fig. 2A).

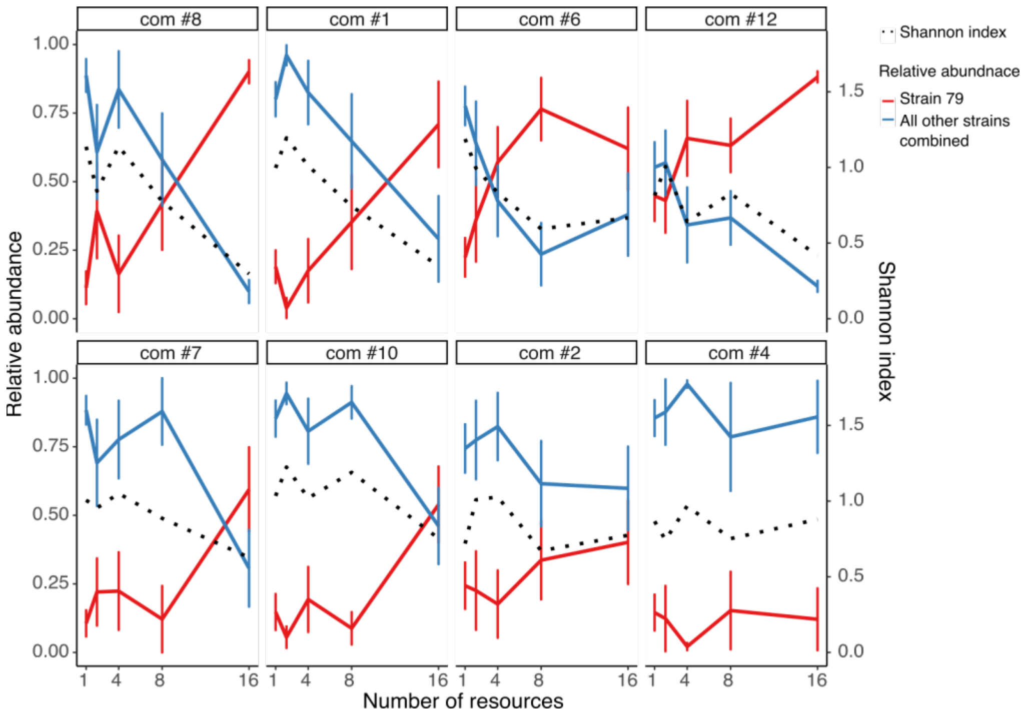

**Supplementary Figure 12: Strain '79' dominance increases with resource complexity, reducing diversity.** Supplementary results for Fig. 2F. Strain '79' exhibits a sharp increase in relative abundance with resource complexity, while the combined relative abundance of all other strains declines. Shannon's diversity index follows this trend, decreasing in parallel with the overall reduction of other strains. Shown are all eight communities containing strain '79', ordered by the slope of Shannon diversity decline between individual resources and 16-resource conditions. The trend line represents the average, with error bars indicating ± SEM.

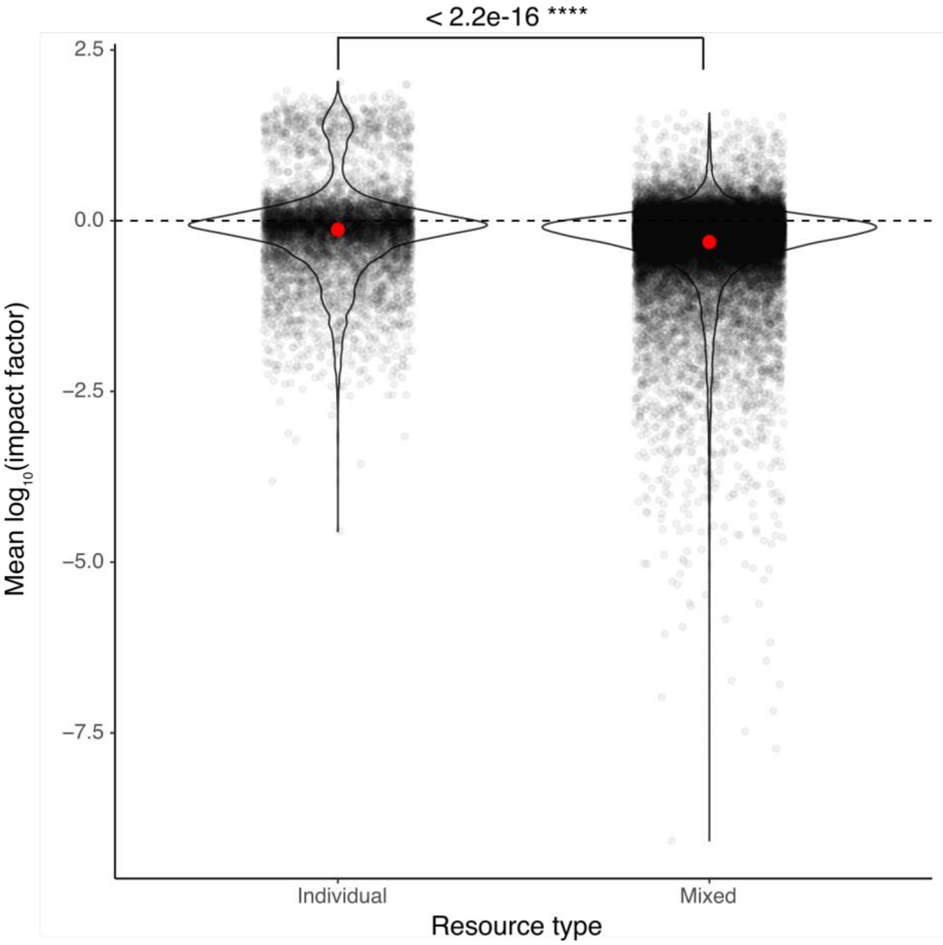

**Supplementary Figure 13: Impact factor on the sentinel strain is more suppressive in mixed resources.** The red dot represents the mean, and the dashed line marks zero impact, indicating no effect on sentinel strain growth. Statistical significance was assessed using a Mann-Whitney U test. In total, 6,556 impact factor values were calculated for individual resources and 26,660 for resource mixtures.

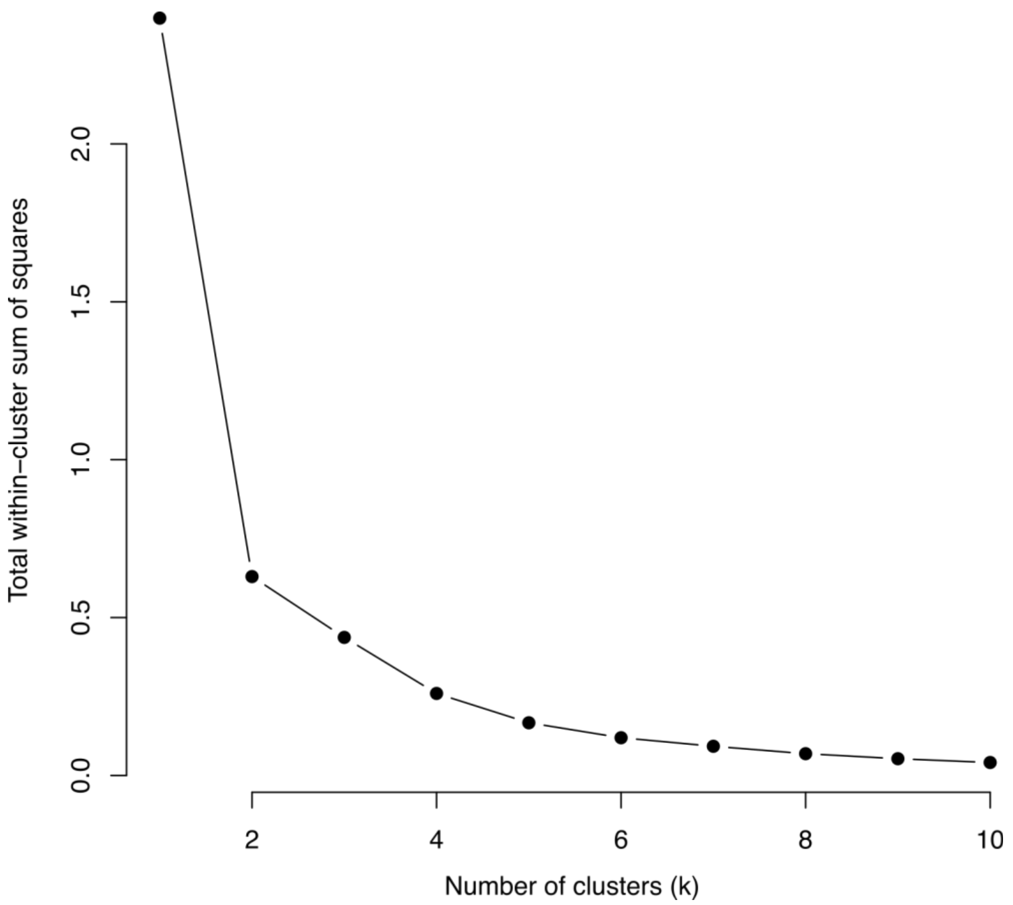

**Supplementary Figure 14: K-means clustering was used to classify strains into two types (highly and mildly influenced).** The optimal number of clusters was determined using the elbow method, where the ‘elbow’ represents the point at which adding more clusters results in only a minimal reduction in clustering error. As shown, the optimal number of clusters is two.

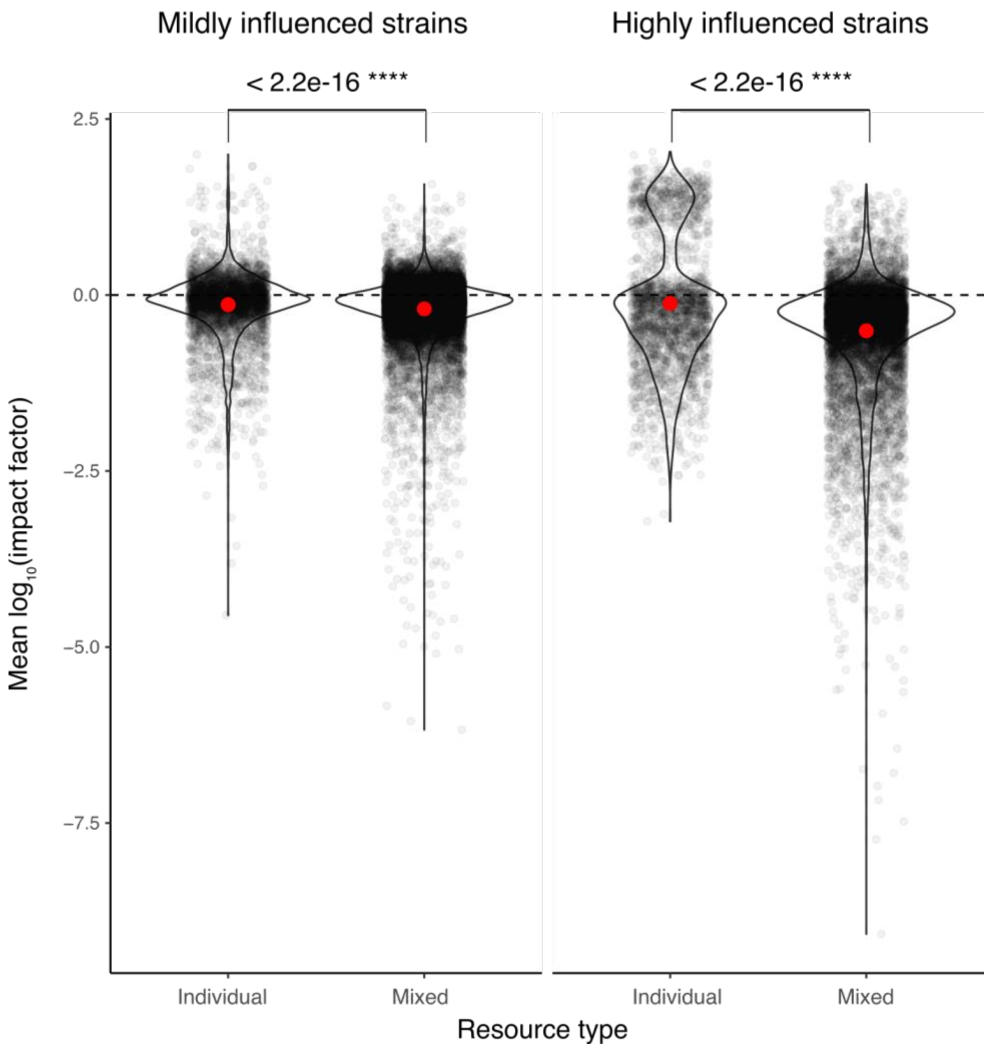

**Supplementary Figure 15: Impact factor on the sentinel strain is more suppressive in mixed resources for both highly and mildly influenced strains.** The red dot represents the mean, and the dashed line marks zero impact, indicating no effect on sentinel strain growth. Statistical significance was assessed using a Mann-Whitney U test. In total, 6,556 impact factor values were calculated for individual resources and 26,660 for resource mixtures.

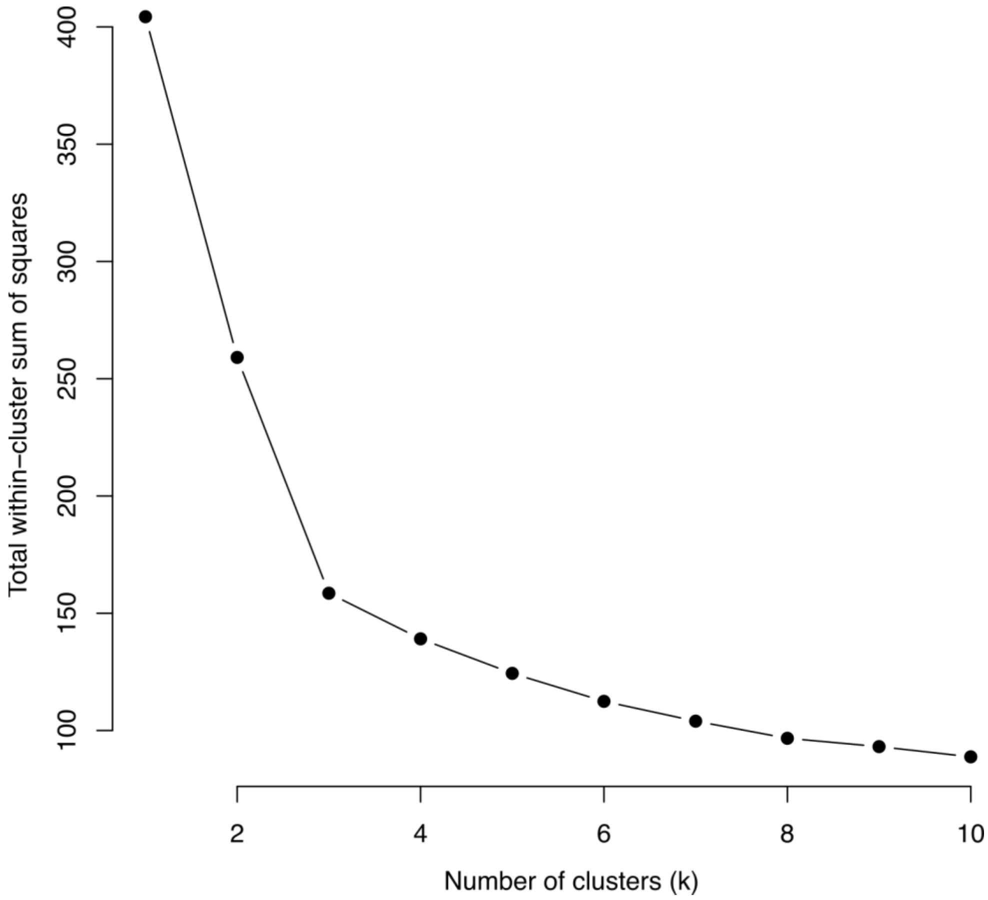

**Supplementary Figure 16: K-means clustering was used to classify strains into three types based on their interactions in individual resources.** The optimal number of clusters was determined using the elbow method, where the ‘elbow’ represents the point at which adding more clusters results in only a minimal reduction in clustering error. As shown, the optimal number of clusters is three.

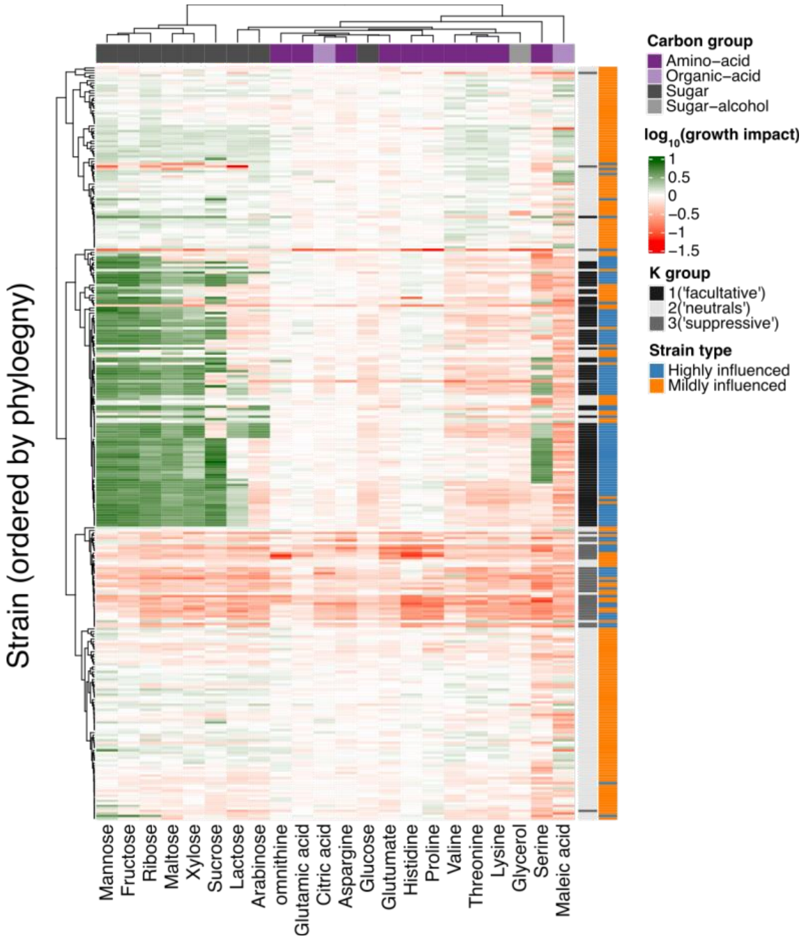

**Supplementary Figure 17: Highly and mildly influenced strains are categorized into three interaction-type clusters based on their behavior in individual resources.** clustering to three groups decided based on supplementary Fig. 16. Highly influenced strains primarily fall into the ‘facultative’ group (K-group 1; shifting from facilitation in sugars to suppression in acids) and the ‘suppressive’ group (K-group 3; consistently suppressive). In contrast, mildly influenced strains mainly belong to the ‘neutral’ group (K-group 2; exerting only mild effects on the pathogen without significantly impacting its growth). The same data is presented in Fig. 3D (‘individual resources’ heatmap).

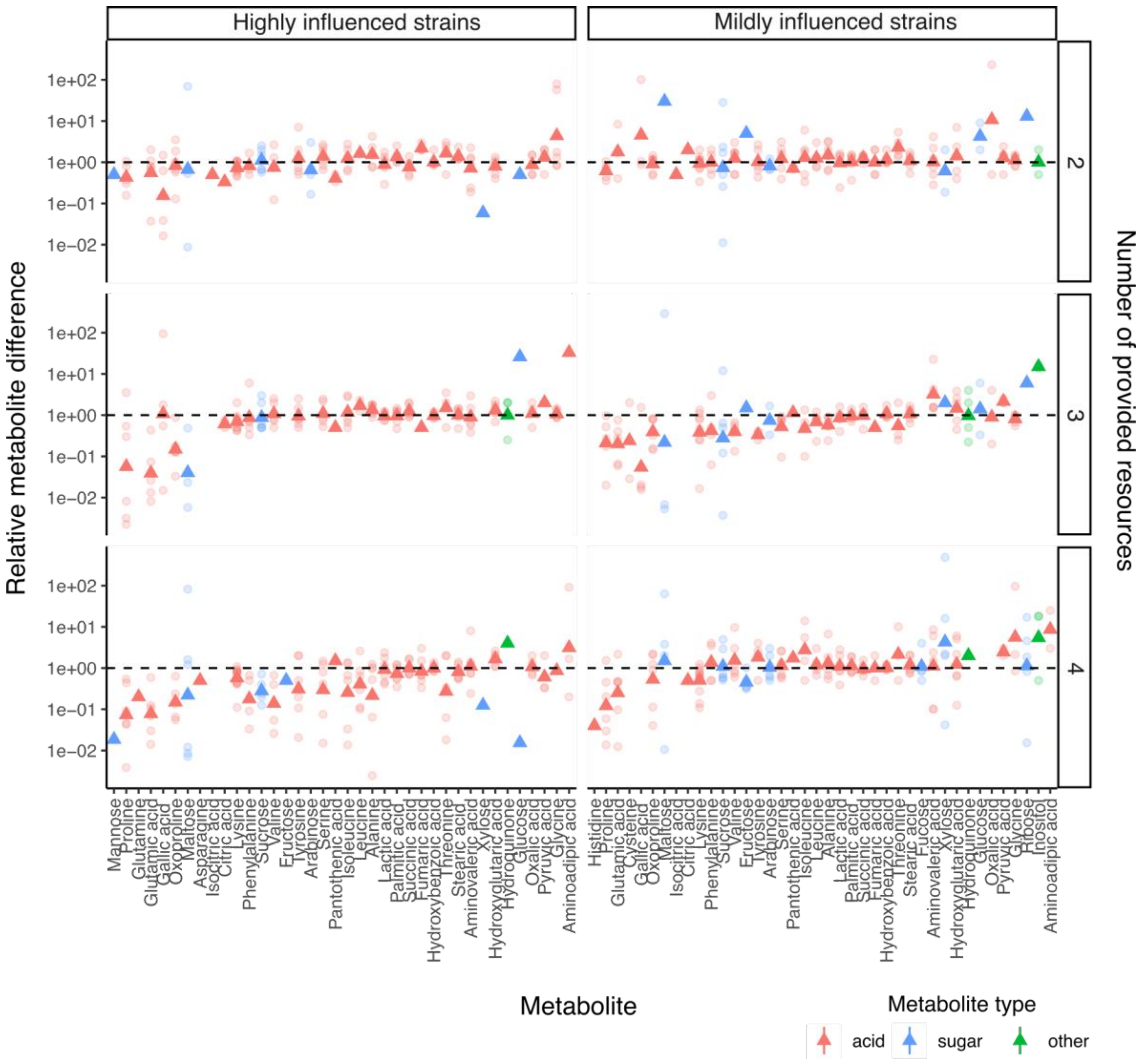

**Supplementary Figure 18: Highly influenced strains show greater metabolic uptake in resource mixtures than in their supernatant mixtures for specific metabolites, with this effect intensifying as resource complexity increases.** Supplementary results for Fig. 4B. Shown is the relative change in metabolite peak area (normalized to an internal control) for 96 identified metabolites, comparing resource mixtures to their matching supernatants. Points represent individual metabolites within a specific sample, while triangles indicate the mean. The y-axis is displayed on a $\log_{10}$ scale.

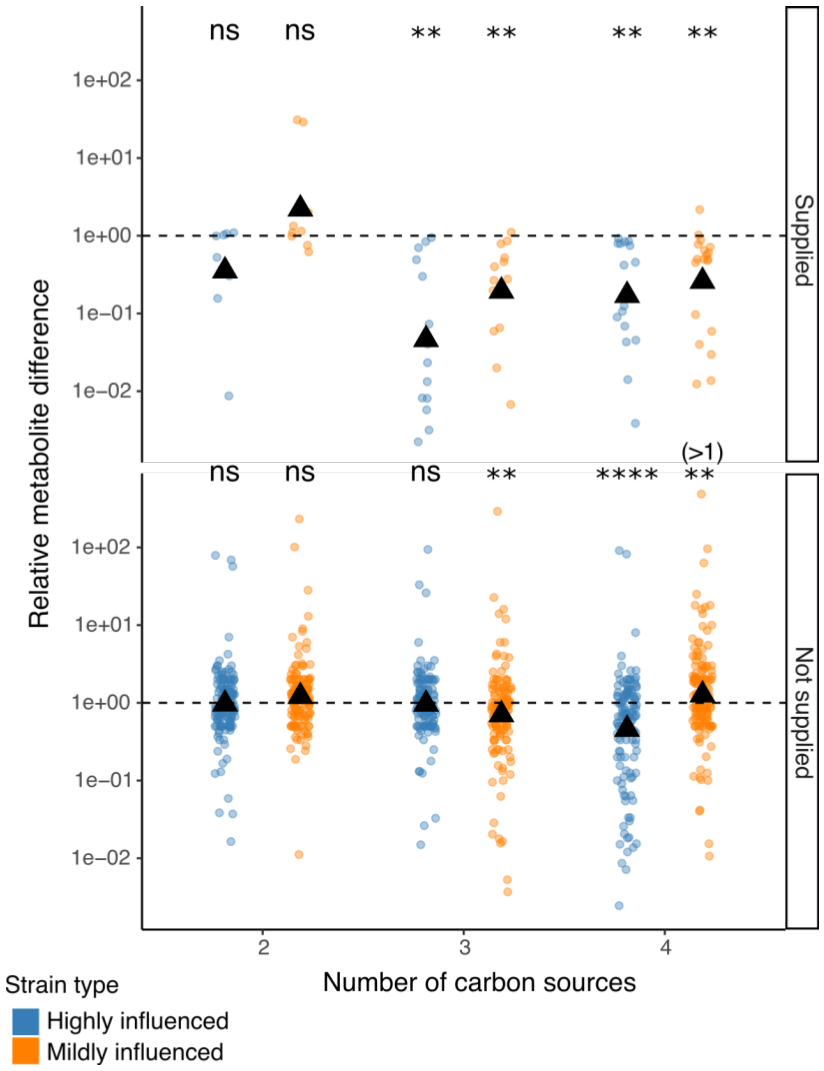

**Supplementary Figure 19: Highly influenced strains show greater metabolic uptake in resource mixtures than in their supernatant mixtures, for both supplied and not supplied resources, with this effect intensifying as resource complexity increases.** Supplementary results for Fig. 4B. Shown is the relative change in metabolite peak area (normalized to an internal control) for 96 identified metabolites, comparing resource mixtures to their matching supernatants. Data are binned by 'supplied' resources (metabolites that were part of the defined media and initially added) and 'not supplied' resources (metabolites likely excreted, as they were not provided in the growth media). Points represent individual metabolites within a specific sample, while triangles indicate the mean. The y-axis is displayed on a $\log_{10}$ scale. A Mann-Whitney U test assessed whether the median relative metabolite difference significantly deviated from 1. For mildly influenced strains with four resources in the 'not supplied' category, values >1 indicate more metabolites remaining in the resource mixture (lower uptake), whereas significance otherwise indicates higher uptake. Significance levels: ns ($p > 0.05$), * ($p \leq 0.05$), ** ($p \leq 0.01$), *** ($p \leq 0.001$), **** ($p \leq 0.0001$).

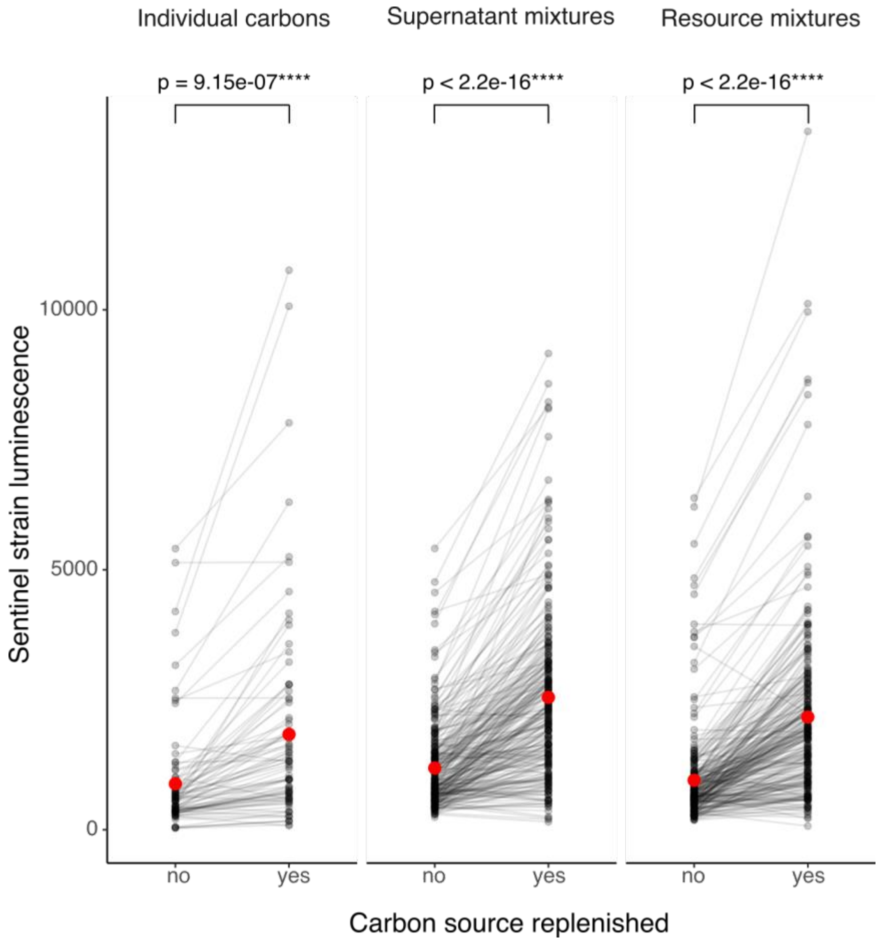

**Supplementary Figure 20: Media replenishment restored normal growth of the sentinel strain in most supernatants, ruling out toxicity as a primary competition driver.** Supplementary results for Fig. 4C. Fresh media containing the matching resources for each sample was added at a 1:10 dilution (9 parts supernatant, 1 part fresh media). The sentinel strain was grown in supernatants with and without replenishment. As shown, replenishment restored growth in most cases. Points represent individual samples (sentinel strain growth in a specific supernatant), with lines connecting the same samples before and after replenishment. The red dot indicates the mean. Replenishment was performed across all three supernatant types—individual carbon sources, supernatant mixtures, and resource mixtures (illustrated in Fig. 4A). Statistical significance was assessed using a Mann-Whitney U test. In total, 84 individual carbon supernatants, 252 supernatant mixtures, and 252 resource mixtures were tested, each with and without replenishment (represented as separate conditions on the x-axis).

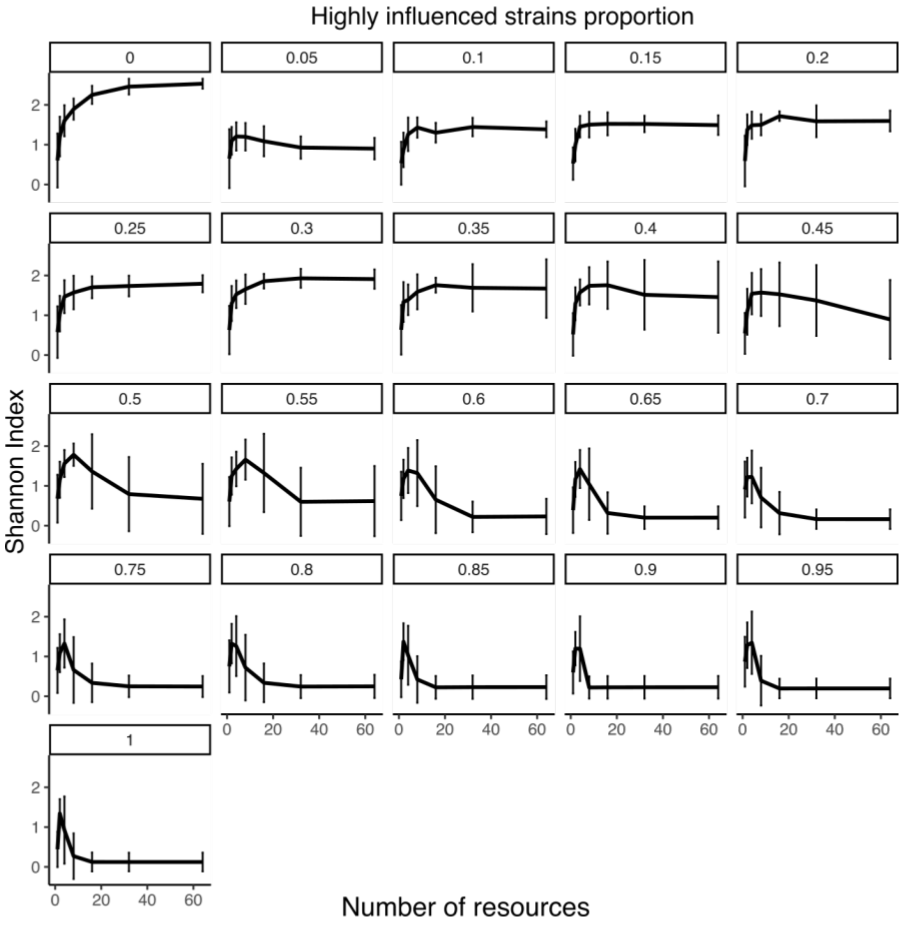

**Supplementary Figure 21: Highly influenced strain behavior reduces microbial diversity as resource complexity increases in a consumer-resource model.** Supplementary results for Fig. 4E. We simulated 420 communities, each with 50 randomly generated strains (assigned random uptake and growth rates for 100 resources) across seven resource complexities (1–64) and 21 highly influenced strain proportions. Specifically, for each proportion (0–100% in 5% steps), 20 communities were simulated. The 0% proportion represents the default consumer-resource model (similar to Fig. 1A). The trend line represents the mean of 20 communities per highly influenced fraction, with error bars indicating ± SD. This analysis used $\gamma = 0.05$.

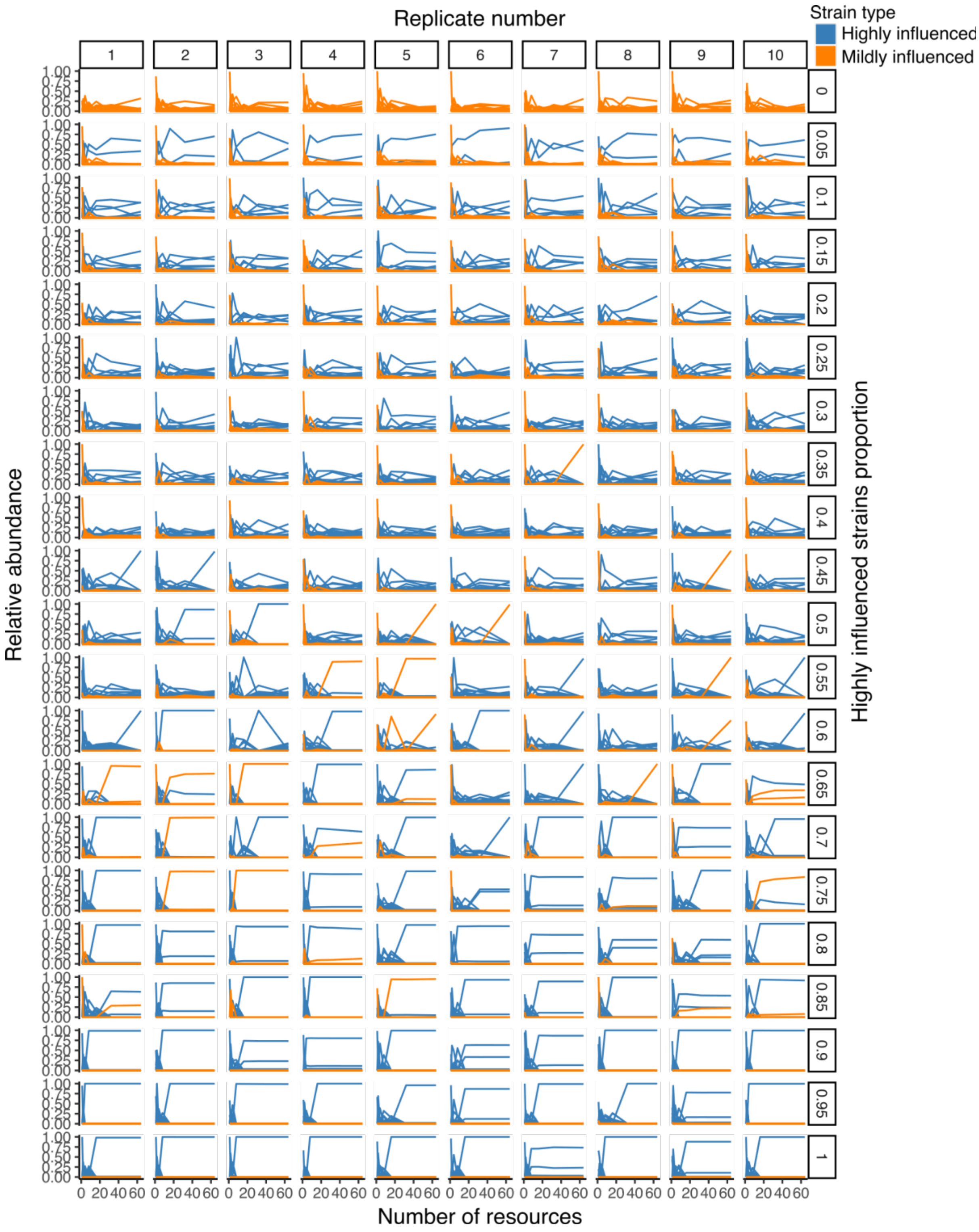

**Supplementary Figure 22: Highly influenced strain behavior reduces microbial diversity as resource complexity increases in a consumer-resource model, with highly influenced strains increasingly dominating.** Supplementary results for Fig. 4E and Supplementary Fig. 14. Shown is the relative abundance of each of the 50 strains per community across highly influenced strain proportions (0–100% in 5% steps), categorized as highly or mildly influenced. Ten randomly selected replicates are displayed (out of 20 total replicates). This analysis used $\gamma = 0.05$.

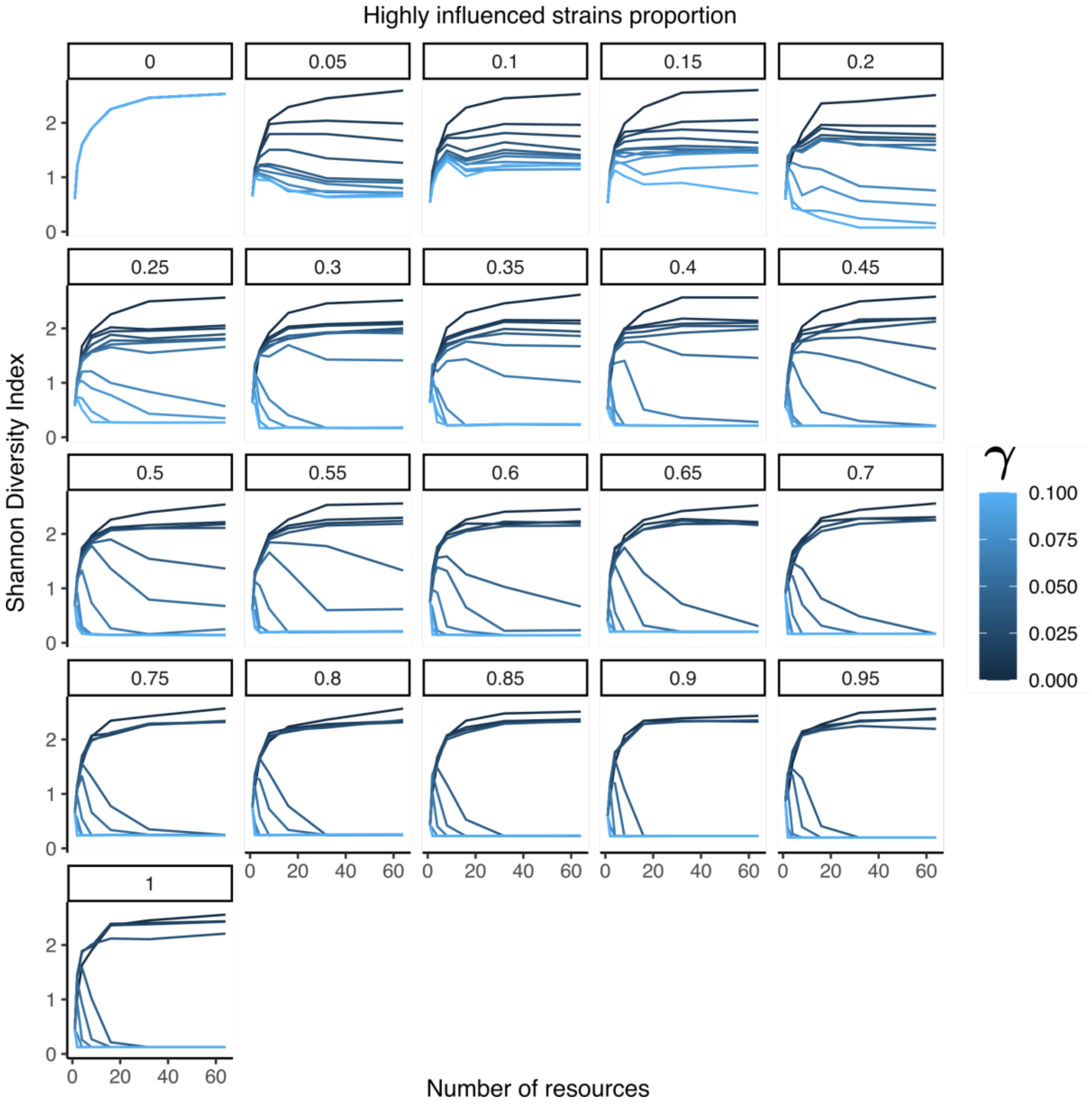

**Supplementary Figure 23: Highly influenced strain behavior reduces microbial diversity as resource complexity increases in a consumer-resource model, with consistent results for γ > 0.03.** Supplementary results for Fig. 4E. Similar to Supplementary Fig. 21, but with a range of γ values (affinity increase factors; Supplementary Text 3) from 0 (default model) to 0.1, increasing in increments of 0.01 (11 γ values in total). The trend line represents the mean of 20 communities per highly influenced strain fraction and γ value.

#### Supplementary Tables

**Supplementary Table 1: Resources used in this study.** A list of carbon compounds categorized by biochemical group, associated metabolic pathway (glycolysis or gluconeogenesis), and vendor information.

| Carbon | Carbon group | Pathway | Vendor |
|---|---|---|---|
| Glucose | Carbohydrate | glycolysis | Sigma-Aldrich |
| Sucrose | Carbohydrate | glycolysis | Sigma-Aldrich |
| Glutamic-acid | Amino-acid | gluconeogenesis | Sigma-Aldrich |
| Glycerol | Sugar-alcohol | both | Sigma-Aldrich |
| Citric acid | Organic-acid | gluconeogenesis | Sigma-Aldrich |
| omnithine | Amino-acid | gluconeogenesis | Sigma-Aldrich |
| Lactose | Carbohydrate | glycolysis | Sigma-Aldrich |
| Maltose | Carbohydrate | glycolysis | Sigma-Aldrich |
| arabinose | Carbohydrate | glycolysis | Sigma-Aldrich |
| Mannose | Carbohydrate | glycolysis | Thermo Fisher Scientific |
| Xylose | Carbohydrate | glycolysis | Sigma-Aldrich |
| Lysine | Amino-acid | gluconeogenesis | Sigma-Aldrich |
| Maleic acid | Organic-acid | gluconeogenesis | Sigma-Aldrich |
| Ribose | Carbohydrate | glycolysis | Sigma-Aldrich |
| Serine | Amino-acid | gluconeogenesis | Merck |
| Proline | Amino-acid | gluconeogenesis | Merck |
| fructose | Carbohydrate | glycolysis | Millipore |
| Threonine | Amino-acid | gluconeogenesis | Merck |
| Valine | Amino-acid | gluconeogenesis | Millipore |
| Histidine | Amino-acid | gluconeogenesis | Merck |
| Aspargine | Amino-acid | gluconeogenesis | Thermo Fisher Scientific |
| Glutumate | Amino-acid | gluconeogenesis | Biowest |

### Supplementary Text

##### Supplementary Text 1: Consumer-resource model

### 1 Classic Consumer-Resource Model

The standard consumer-resource model follows Michaelis-Menten kinetics for resource uptake, with species growth determined by available resources.

#### 1.1 Species Dynamics (Classic)

The change in abundance of species $i$ over time follows:

$$\frac{dN_i}{dt} = N_i \min\left(\sum_\alpha e_{i\alpha} v_{0,i\alpha} \frac{R_\alpha}{km_{i,\alpha} + R_\alpha},\ r_{0,i}\right) - \text{dil} \cdot N_i. \tag{1}$$

where:

- $N_i$ is the abundance of species $i$,
- $r_{0,i}$ is the maximum intrinsic growth rate of species $i$,
- $v_{0,i\alpha}$ is the maximum uptake rate of resource $\alpha$ by species $i$,
- $km_{i,\alpha}$ is the half-saturation constant governing affinity for resource $\alpha$,
- $R_\alpha$ is the concentration of resource $\alpha$,
- $e_{i\alpha}$ is the efficiency of converting resource $\alpha$ into biomass,
- dil represents the dilution rate (washout in a chemostat-like system).

Essentially, we assume each resource follows its own Michaelis-Menten kinetics and contributes to growth rate additively. When resources are so abundant that they no longer limit growth rate, the growth rate takes the maximum intrinsic value $r_{0,i}$.

#### 1.2 Resource Dynamics (Classic)

The change in resource concentration follows:

$$\frac{dR_\alpha}{dt} = -\sum_i (c_{i\alpha} N_i) + \sum_{i,\beta} b_{i,\beta\to\alpha}(1 - e_{i\beta}) c_{i\beta} N_i - \text{dil} \cdot R_\alpha + \text{dil} \cdot \text{Rsupp}_\alpha. \tag{2}$$

where:

- $c_{i\alpha}$ is the actual consumption rate, given by:

  $$c_{i\alpha} = v_{0,i\alpha} \frac{R_\alpha}{km_{i,\alpha} + R_\alpha} \times \frac{\min(a_i, r_{0,i})}{a_i}$$

  In other words, we assume that when resources are abundant and growth is limited by other factors, the species scale down the uptake of each resource proportionally without preference.

- $b_{i,\beta\to\alpha}$ represents the fraction of resource $\beta$ secreted as resource $\alpha$, and satisfies $\sum_\alpha b_{i,\beta\to\alpha} \leq 1$.

**Supplementary Text 2: Inferring interactions matrices from experimental data using the generalized Lotka-Volterra model**

### 1 Data and Methodology

Data from single-species growth, pairwise interactions (Fig. 2 A) and community composition (Fig. 1 D, E and Supplementary Fig. 3) at steady state was used to infer the interaction matrices of the 14 strains comprising the experimental communities. The dataset was divided into three groups based on carbon source richness: 1, 8, and 16 resources. Each group contained multiple nutrient combinations, so we did not assume a fixed interaction strength for a given strain pair across conditions. Instead, we treated interaction strengths as random variables drawn from an unknown distribution and used Bayesian regression to infer their values.

### 2 Generalized Lotka–Volterra Model at Steady State

With the generalized Lotka–Volterra (gLV) model we obtain under steady-state assumptions:

$$\frac{dx}{dt} = k \circ x \circ (1 - Ax) = 0, \tag{1}$$

where $x$ is an $N$-dimensional vector containing steady-state population densities of N interacting species, $A$ is the $N \times N$ interaction matrix, k a N-dimensional vector of growth rates, and $\circ$ the Hadamard product (element-wise multiplication). We take into account only the species that are present in the community in steady state ($x_i > 0$). For these species the equation (1) can only become zero if $(1 - Ax)$ equals zero. Accordingly, equation (1) simplifies to:

$$\mathbf{1}_{\{x_i>0\}} \circ (1 - Ax) = 0, \tag{2}$$

with $\mathbf{1}_{\{x_i>0\}}$ denoting a N-dimensional indicator vector, that has entry 1 if the species i is present at steady state and 0 otherwise.

### 3 Bayesian Regression and MCMC Implementation

Equation (2) was fit using Bayesian regression with Monte Carlo Markov Chain (MCMC) sampling, implemented via the NUTS solver in the Python `pyro` package.

#### 3.1 Priors and Likelihood

Priors for the off-diagonal elements ($\alpha_{\text{other}}$) of the interaction matrix A were assumed to be normally distributed with mean 1 and variance 1. Priors for the diagonal elements ($\alpha_{\text{self}}$) of the interaction matrix A were normally distributed with mean equal to the inverse of the carrying capacity (i.e., the steady-state $OD_{600}$ in monoculture, Supplementary Fig. 7) and variance $5 \times 10^{-6}$. With

$$y = \mathbf{1}_{\{x_i>0\}} \circ (1 - Ax) \tag{3}$$

the likelihood for the Bayesian regression was defined as:

$$\mathcal{L}(y|x, A) \sim \mathcal{N}(0, 10^{-5}\mathbf{I}). \tag{4}$$

#### 3.2 Posterior Sampling

Posterior sampling was performed using 30 chains, each with 400 warm-up steps and 200 sampling steps. Convergence was assessed by plotting posterior traces over time, and the final interaction matrix values were computed as the mean of the posterior samples. The software code for the Bayesian Regression can be accessed at https://github.com/RatzkeLab .

**Supplementary Text 3: Consumer-resource model with highly influenced strains modification**

### 1 Highly Influenced Strains Modification: Dynamic $km_{i,\alpha}$

We observed that highly influenced strains use resources more thoroughly than mildly influenced strains as resource number increases (Fig. 4B). We incorporated this behavior in the highly influenced strains modified version as follows: species' half-saturation constants ($km_{i,\alpha}$) dynamically decrease as resource diversity increases.

#### 1.1 Highly Influenced Strains Modified Half-Saturation Constant

For highly influenced strains ($\gamma_i > 0$):

$$km_{i,\alpha} = \frac{km^0_{i,\alpha}}{1 + \gamma_i \sum_\beta \mathbf{1}_{\{R_\beta > \theta_R\}}}. \tag{1}$$

For mildly influenced strains ($\gamma_i = 0$), we retain the classic model:

$$km_{i,\alpha} = km^0_{i,\alpha}. \tag{2}$$

where:

- $km^0_{i,\alpha}$ is the baseline half-saturation constant for species $i$ and resource $\alpha$.
- $\gamma_i$ is a species-specific coefficient controlling adaptation to resource diversity.
- $\sum_\beta \mathbf{1}_{\{R_\beta > \theta_R\}}$ counts how many resources exceed a small availability threshold $\theta_R$, where $\mathbf{1}_{\{R_\beta > \theta_R\}}$ is an indicator function.

#### 1.2 Highly Influenced Strains Modified Species Dynamics

Replacing $km_{i,\alpha}$ in Equation (1):

$$\frac{dN_i}{dt} = N_i \min\left(\sum_\alpha e_{i\alpha} v_{0,i\alpha} \frac{R_\alpha}{\frac{km_{0,i,\alpha}}{1+\gamma_i \sum_\beta \mathbf{1}(R_\beta > \text{threshold})} + R_\alpha},\ r_{0,i}\right) - \text{dil} \cdot N_i. \tag{3}$$

#### 1.3 Highly Influenced Strains Modified Resource Dynamics

Substituting $km_{i,\alpha}$ in Equation (2):

$$\frac{dR_\alpha}{dt} = -\sum_i (c_{i\alpha} N_i) + \sum_{i,\beta} b_{i,\beta\to\alpha}(1 - e_{i\beta}) v_{0,i\beta} \frac{R_\beta}{\frac{km_{0,i,\beta}}{1+\gamma_i \sum_\gamma \mathbf{1}(R_\gamma > \text{threshold})} + R_\beta} N_i - \text{dil} \cdot R_\alpha + \text{dil} \cdot \text{Rsupp}_\alpha. \tag{4}$$

#### 2 Key Differences from the Classic Model

- In the **classic model**, $km_{i,\alpha}$ is fixed for each species-resource interaction.
- In the **highly influenced strains modified model**, $km_{i,\alpha}$ decreases dynamically when multiple resources exceed the threshold, improving uptake efficiency.
- This effect is controlled by the parameter $\gamma_i$:
  - If $\gamma_i = 0$ (mildly influenced strains), the species behaves like in the classic model.
  - If $\gamma_i > 0$ (highly influenced strains), the species adapts its uptake based on resource diversity.

#### 3 Parameter Values

By default we set $e \equiv 1$ so that resources $R$ are fully converted to biomass $N$, and there is no cross-feeding. We also set $v \equiv 1$ so the maximum uptake rate is the same for different resources.

#### 4 Conclusion

This model extends the classic consumer-resource framework by introducing a resource-diversity-dependent adjustment to uptake affinity ($km$). This modification enables species to respond dynamically to resource availability, which may lead to competitive advantages in diverse environments.
